# Linking Genomic Landscape to Disease Mechanism: Core Genetic Factors Underlying Pathogenesis and Antimicrobial Resistance in Diarrheal Pathogens

**DOI:** 10.1101/2025.08.31.673274

**Authors:** Mohammad Uzzal Hossain, Marjia Akter Suchi, Zeba Sanjida, A.B.Z. Naimur Rahman, Mohammad Nazmus Sakib, SM Sajid Hasan, Md. Ekram Hossan, Abu Nayeem Ahmed, Mahmudul Hasan, Tamal Paul, Arittra Bhattacharjee, Zeshan Mahmud Chowdhury, Ishtiaque Ahammad, M.M. Kamal Hossain, Palash Kumar Sarker, Farha Matin Juliana, Md. Salimullah, Keshob Chandra Das

## Abstract

**Background:** Diarrheal diseases remain a major global health burden, as they severely affect children, particularly in Bangladesh. After decades of research, the molecular mechanisms of diarrheal pathogens for disease pathogenesis and antibiotic resistance are still unknown, notably in Gram-negative bacteria. This pilot study fills the gap by employing whole genome sequencing and pan-genome analysis to analyze Bangladeshi diarrheal pathogens to identify genetic variables that cause disease pathogenesis and antibiotic resistance.

**Results:** Hence, we investigated the genetic diversity of bacterial isolates from 31 clinical stool samples by a combination of whole-genome sequencing (WGS) and pan-genomic analysis. A core group of 50 genes, conserved across a significant number of strains, was identified via pan-genomic analysis, with considerable variation in accessory genes. This signifies a significant degree of genetic flexibility. Gene ontology analysis yielded substantial insights into prospective therapeutic targets by emphasizing the critical function of these core genes in bacterial survival and pathogenicity. Furthermore, the findings of the antimicrobial susceptibility test (AST) revealed concerning resistance trends, particularly to fluoroquinolones and beta-lactams, underscoring the necessity for enhanced surveillance and alternative therapeutic approaches.

**Conclusion:** This study provides a comprehensive genetic framework to improve understanding of the complexity of diarrheal infections and the mechanisms underlying their resistance, fostering opportunities for potential therapeutic advancements.

## Background

Diarrhoea is defined as the expulsion of three or more loose or liquid stools within a day, or a frequency exceeding an individual’s normal pattern. It is a global problem that affects all regions and populations (1,2). Diarrheal diseases are a significant cause of illness and death among children under five, leading to approximately 500,000 deaths each year. It is the third largest contributor to the global burden of disease as assessed by disability-adjusted life years (DALYs) (3,4). Diarrhoea has the biggest burden in low- and middle-income countries (LMICs), where it is responsible for nearly 20% of deaths among children aged 1 to 59 months in regions like Sub-Saharan Africa and South Asia (4). In Bangladesh, diarrheal disease is one of the leading causes of death and the most common reason for hospitalization. (5–7). Research have indicated that the overall prevalence of diarrhoea among children under the age of five in Bangladesh stands at 5.71% (8).

Although Diarrhoea can be caused by a number of factors, infectious agents such as bacteria and viruses are the most frequent. For bacteria, gram-negative enteric bacterial pathogens (GNEBPs) most significantly contribute in illness and predominantly located in the intestine. These pathogens come from different genera including *Escherichia, Shigella, Campylobacter, Salmonella, Enterobacter, Klebsiella, Yersinia, Serratia,* and *Proteus*, among others. However, the most prevalent and major pathogens include *S. enterica, E. coli, Campylobacter*, and *Shigella spp*. (9).

Previous studies in Bangladesh highlighted gram-negative bacteria as major players (8) in diarrheal diseases. Among these bacteria, *Shigella flexneri* is the most commonly identified (54%-60%) gastrointestinal pathogen (10) and it is estimated that more than 95,000 children under the age of five die each year from shigellosis (11,12). *S. flexneri* is extremely contagious, and 10^2^ - 10^3^ bacteria are sufficient to cause diarrhoea in humans (13,14). Additionally, *Vibrio cholerae* is a common cause of diarrheal epidemics and outbreaks throughout Bangladesh. Despite the efforts to prevent Diarrhoeal diseases over the last few decades, recent data suggests sporadic outbreaks in several districts of Bangladesh, resulting in a large number of cases and deaths (15). While these studies have provided valuable insights into the prevalence and impact of gram-negative bacterial pathogens, there still remains a gap in understanding the genetic diversity and evolutionary dynamics of these pathogens. This gap can be addressed through pangenome analysis to characterize the genome, facilitating to identify the core, accessory and shell genes. Moreover, the antibiotic resistance profiles of these isolated gram-negative bacteria can enhance the understanding of the resistance mechanism.

In this study, we performed a comprehensive pangenome analysis of gram-negative bacterial pathogens isolated from diarrheal patient in Bangladesh. Whole-genome sequencing served as the foundation for this powerful strategy, enabling us to identify core and accessory genes shared among closely related microbes through pan-genome. Pan-genome represents the complete set of genes found across a collection of isolates (e.g. strains from one species). It helps capture the diversity and dynamics of genomes within a specific taxonomic group, whereas individual genomes often only represent a small portion of the pangenome (16,17). Three methods of pan-genome analysis are frequently employed to examine the genetic makeup of microorganisms: core genome profiling, accessory genome profiling, and specific genome profiling. These methods demonstrate the traits of homology, variety, and specialization among genomes (18,19). For example, the core genome, which consists of genes shared by all individuals, is frequently used to determine the relatedness of strains within the same species (20,21). Furthermore, pangenome research indicated considerable horizontal gene transfer within the accessory genomes of microorganisms (22). Pan-genome analysis has been used to examine the genetic diversity of related microbial genomes, providing insights into gene composition, strain tracking, evolutionary impacts, niche specialization, screening for antimicrobial targets, and discovering diagnostic markers (23–27). Additionally, we conducted antibiotic resistance profiling of these gram-negative bacteria, which may facilitate targeted treatment, avoid resistance, cut global healthcare expenditures, and significantly contribute to saving lives. This knowledge assists healthcare personnel in selecting the most effective medicines for a patient’s unique medical condition, preventing the overuse of broad-spectrum antibiotics and enhancing patient outcomes.

With the advancements of Next-Generation Sequencing (NGS), WGS, extensive bioinformatics tools and systems biology approaches, performing pan-genome analysis has become significantly easier and less complex. NGS enables high-throughput sequencing, providing vast amounts of genomic data that can be analyzed with precision and efficiency. Whole-genome sequencing (WGS) and phylogenomics have considerably improved pathogen surveillance and allow for more thorough monitoring. The dynamics of pathogenic bacterial genomes are poorly known, particularly in LMICs such as Bangladesh. The absence of comparable research in Bangladesh, where gram-negative bacteria-associated diarrhoea poses a considerable health burden, emphasizes the urgent need for a complete understanding of genetic profiles, epidemiology, resistance and dynamics.

At Hossain *et al*. lab, three significant studies were performed to investigate *Plesiomonas shigelloides* (28)*, Providencia stuartii* (29) and *Citrobacter werkmanii* (30) isolated from diarrheal patient. In these study, next-generation sequencing, pangenome analysis, system biology approaches were applied to explore comparative genomic analysis and coinfection pattern among diarrheal pathogens. Therefore, we utilized the same methodology to conduct our experiments and extends these analyses to further explore the genomic landscape to characterize the genetic insights of gram-negative diarrheal pathogens in Bangladesh.

## Methods

### Ethical approval

The approval for sample collection and laboratory experiments was received from the Institutional Review Board (IRB) under Approval No: NIB/IRB/2023/BID-04. Ethical standards were maintained accordingly and every participant who were part of the research had to provide their consent. The consent was collected either verbally or in writing.

### Clinical Sample Collection

31 stool samples were collected from children (age ≥ 8) suffering from diarrhoea at Dhaka Sishu Hospital, Dhaka (23.7732° N, 90.3628° E), and Lakshmipur Sadar Hospital, Lakshmipur (22.934737°N, 90.832875°E). The sample metadata presented in **Supplementary Table S1** includes demographic information such as age, geographic origin, and gender of the patients. Using sterile specimen containers, the samples were stored in Amies medium for transport and then quickly transported to laboratory within 2-3 hours for further analysis including isolation and identification studies.

### Isolation and Identification of Bacteria

Samples were inoculated into MacConkey agar (Oxoid, UK) plate and incubated at 37°C for 24 hours. All suspected colonies from the selective media were then isolated, streaked onto nutrient agar plates, and incubated at 37°C for 18 to 24 hours and typical colonies from nutrient agar plates were selected and subjected to slide preparation, gram staining, and microscopic examination. Biochemical identification was carried out using the VITEK 2 compact system (bioMérieux, France).

### Antibiotic Susceptibility Test

Antimicrobial susceptibility was analysed based on the Kirby disc diffusion method (31). The susceptibility of the isolate was evaluated using antibiotic disks: Ampicillin (AMP, 10 μg), Cefotaxime (CTX, 30 μg), Ceftazidime (CAZ, 30 μg), Cefepime (FEP, 30 μg), Gentamycin (CN, 10 μg), Streptomycin (S, 10 μg), Tetracycline (TE, 30 μg), Chloramphenicol (C, 30 μg), Ciprofloxacin (CIP, 5 μg), Levofloxacin (LEV, 5 μg), Nalidixic Acid (NA, 30 μg), Sulfamethoxazole/Trimethoprim (SXT, 25 μg), Meropenem (MEM, 10 μg), Imipenem (IMI, 10 μg) and Azithromycin (AZM, 15 μg). All antibiotic disks were sourced from Oxoid Ltd, UK. Zones of inhibition were measured for each antibiotic and interpreted as sensitive, intermediate, or resistant according to the Clinical and Laboratory Standards Institute (CLSI) guidelines (32).

### Bacterial Genomic DNA Extraction and Purification

Genomic DNA of bacterial isolates was extracted and purified using the Thermo Scientific GeneJET Genomic DNA Purification Kit, following the manufacturer’s extraction protocol. This kit employs silica-based membrane technology in the form of a spin column, allowing for rapid and efficient isolation of high-quality genomic DNA from bacterial samples. After the purification process, DNA concentration and purity were assessed using a NanoDrop™ 2000/2000c Spectrophotometer to ensure the quality of the extracted genomic DNA.

### Whole-genome sequencing and bioinformatic analysis

The confirmed bacterial isolates were sequenced using the Illumina MiSeq platform at the National Institute of Biotechnology (NIB), which provides next-generation sequencing services. DNA libraries were constructed according to the manufacturer’s instructions using the Illumina Nextera XT DNA Library LT kit (San Diego, CA, USA) and whole-genome sequencing (WGS) was carried out on the Illumina MiSeq system with a 300 bp paired-end read sequencing kit (MiSeq Reagent Kit v3; Illumina).

After sequencing, the raw data from the MiSeq instrument in the gz compressed FASTQ format were first assessed for quality control using the Linux terminal and the FastQC (33) command tools to evaluate read quality, GC content, and potential contamination. Then, low-quality bases and adapter sequences were trimmed using software such as Trimmomatic (34). *De novo* genome assembly was performed using Unicycler (35) which employs Spades (version 3.15.5) with error correction to minimize mismatching to generate a De Bruijn graph assembly by analysing a diverse range of k-mer sizes with error correction to minimize mismatching. After assembly, the quality of the assembled genome was assessed using the QUAST tool (36). To decipher the genetic content and functional elements encoded within the assembled contigs, the Prokka pipeline (version 1.14.6) (37) was employed for annotation. Genomes were filtered by the number of contigs, assembly size, GC content and predicted number of coding genes.

### Pangenome Analysis Using Roary

69 bacterial WGS sequencing data from clinical samples were retrieved from the National Center for Biotechnology Information (NCBI). The obtained genomes were annotated using the Prokka pipeline (37). The roary tool (38) was used to conduct pan-genome analysis of 100 genome sequences of 10 different bacterial clinical isolates (10 *Citrobacter werkmanii*, 5 *Enterobacter chuandaensis*, 21 *Escherichia coli*, 1 *Escherichia fergusonii*, 13 *Enterobacter hormaechei*, 14 *Klebsiella pneumoniae*, 9 *Morganella morganii*, 11 *Proteus mirabilis*, 6 *Plesiomonas shigelloids*, 10 *Providencia stuartii*) which clustered annotated genes into orthologous groups. This software used GFF3 files as input and generated a presence/absence matrix, core genome alignment, and accessory gene data. Then FastTree (39) was used to construct a midpoint rooting phylogenetic tree and visualized using FigTree. After the identification of core, accessory and unique genes, a python script, ‘roaryplots.py’ (https://github.com/sanger-pathogens/Roary/tree/master/contrib/roary_plots) was utilized to observe the graphs generated as output. Then, Gene ontology function was analysed using tools such as STRING, UniProt, KEGG pathway and RepSeq data (40–43).

### Genomic Feature determination

For the identification of virulence and antimicrobial resistance (AMR) genes, we used ABricate (44), a bioinformatics tool along with two specific databases. The Virulence Factor Database (VFDB) (45) was used for detecting virulence genes associated with diarrheal pathogenicity and the Comprehensive Antibiotic Resistance Database (CARD) (46) was used for identifying genes resistance to antimicrobial agents. The analysis was performed using default parameters, with gene presence determined based on sequence similarity thresholds of ≥80% identity and ≥70% coverage. To identify non-homologous genes within the core genome, we employed BLASTp for sequence alignment and CD-HIT (47) for clustering similar protein sequences based on a defined sequence identity threshold. Additionally, to detect potential toxin-producing genes, we utilized the ToxFinder (https://cge.food.dtu.dk/services/ToxFinder/) tool, which enables the rapid identification of toxin-associated gene sequences within bacterial genomes.

### Pathway Annotation Network determination

We used ClueGO (48) plugin from CytoScape to analyse the 50 core genes in the previous step. This tool helps to differentiate cluster of genes based on functionality. During the analysis, we loaded marker lists of *Escherichia Coli*. Further, we selected GO ontologies, Benjamini-Hochberg correction, and Prefused force directed layout. Benjamini-Hochberg controls the False Discovery Rate (FDR) (49) which is very significant because the expected proportion of false positives can be identified.

## Results

### Initial characterization of the isolates provided insights on 10 unique bacterial strains

Our isolated 31 clinical bacterial samples were gram-negative, rod or comma shaped indicating the typical feature of Enterobacteriaceae family **(Table 1)**. Additionally, mucoid and swarming colonies of *Klebsiella pneumoniae* and *Proteus mirabilis* colonies were identified and lactose and non-lactose fermenters showed their respective characteristics on MacConkey agar plates. MALDI-TOF MS (matrix-assisted laser desorption ionization–time of flight mass spectrometry) revealed 10 unique bacterial strains **(Supplementary Table S2**).

**Table 1:** Frequency distribution of identified bacterial isolates (n=31)

| **Bacterial Species** | **Frequency (n)** | **Percentage (%)** |
| --- | --- | --- |
| *Escherichia coli* | 18 | 58.06 |
| *Klebsiella pneumoniae* | 4 | 12.9 |
| *Enterobacter hormaechei* | 2 | 6.45 |
| *Morganella morganii* | 1 | 3.23 |
| *Proteus mirabilis* | 1 | 3.23 |
| *Escherichia chuandaensis* | 1 | 3.23 |
| *Plesiomonas shigelloides* | 1 | 3.23 |
| *Citrobacter werkmanii* | 1 | 3.23 |
| *Providencia stuartii* | 1 | 3.23 |
| *Escherichia fergusonii* | 1 | 3.23 |
| **Total** | **31** | **100.00%** |

### Metabolic properties were determined by VITEK 2 system

Although bacterial isolates were initially confirmed using MALDI-TOF MS, phenotypic conformation was determined by biochemical tests. 47 tests relevant for Gram-negative identification were compiled (**Table 2**) from the VITEK® 2 system providing detailed understanding on metabolic properties of the isolates although the system contains 64 wells for targeting various biochemical reactions.

**Table 2:** Determination of metabolic properties by VITEK 2 system.

| **Test** | *Klebsiella pneumoniae* | *Morganella morganii* | *Proteus mirabilis* | *Enterobacter hormaechei* | *Escherichia fergusonii* | *Enterobacter chuandaensis* | *Escherichia coli* | *Plesiomonas shigelloides* | *Citrobacter werkmanii* | *Providencia stuartii* |
| --- | --- | --- | --- | --- | --- | --- | --- | --- | --- | --- |
| APPA | - | - | - | - | - | - | - | - | - | - |
| ADO | + | - | - | - | + | - | - | - | - | - |
| PyrA | + | - | - | - | + | - | - | + | + | - |
| IARL | - | - | - | - | - | - | - | - | - | - |
| dCEL | + | - | - | + | + | + | - | - | - | + |
| BGAL | + | - | - | + | + | + | + | + | + | + |
| H<sub>2</sub>S | - | + | + | - | - | - | - | + | + | - |
| BNAG | - | - | - | + | - | + | - | - | - | + |
| AGLTp | - | - | - | - | - | - | - | - | - | - |
| dGLU | + | + | + | + | + | + | + | + | + | + |
| GGT | + | - | - | + | - | + | - | - | + | + |
| OFF | + | + | - | + | + | + | + | + | + | + |
| BGLU | + | - | - | - | - | + | - | - | - | - |
| dMAL | + | - | - | + | + | + | + | + | + | + |
| dMAN | + | - | - | + | + | + | + | + | + | + |
| dMNE | + | - | - | + | + | + | + | + | + | + |
| BXYL | + | - | - | + | - | + | - | - | - | + |
| BAlap | - | - | - | - | - | - | - | - | - | - |
| ProA | + | - | - | - | - | + | - | - | - | - |
| LIP | - | - | - | - | - | - | - | - | - | - |
| PLE | + | - | - | + | - | + | - | - | - | + |
| TyrA | + | - | - | + | - | + | - | - | - | + |
| URE | - | + | + | - | - | - | - | - | - | - |
| dSOR | + | - | - | + | - | + | + | + | + | + |
| SAC | + | - | - | + | - | + | + | - | - | + |
| dTAG | + | - | - | - | + | - | - | - | - | - |
| dTRE | + | + | + | + | + | + | + | + | + | + |
| CIT | + | - | - | + | - | + | - | + | + | + |
| MNT | + | - | - | + | - | - | - | + | + | + |
| 5KG | - | - | - | - | - | - | + | + | + | - |
| ILATk | + | - | - | - | + | + | - | - | + | + |
| AGLU | - | - | - | - | - | - | - | - | - | - |
| SUCT | + | - | - | + | - | + | + | + | + | + |
| NAGA | - | - | - | + | - | + | - | - | - | + |
| AGAL | + | - | - | + | - | - | - | - | - | + |
| PHOS | + | + | + | - | - | - | - | - | - | - |
| GlyA | - | - | - | - | - | + | - | - | - | - |
| ODC | - | - | - | - | + | + | + | - | - | - |
| LDC | + | - | - | - | + | - | + | - | - | - |
| IHISa | - | - | - | - | - | - | - | - | - | - |
| CMT | + | + | + | + | + | + | + | - | - | - |
| BGUR | - | - | - | - | - | - | - | - | - | - |
| O129R | + | - | + | + | - | - | + | - | - | + |
| GGAA | - | - | - | - | - | - | - | - | - | - |
| IMLTa | - | - | - | - | - | - | - | - | - | - |
| ELLM | - | - | - | - | - | - | - | - | - | - |
| ILATa | + | - | - | - | - | + | - | - | - | - |

### Antibiotic susceptibility

Resistance profiles were built upon the results directed from the antibiotic susceptibility testing by following the standard disk diffusion technique (Kirby-Bauer test) from Clinical and Laboratory Standards Institute with interpretations of susceptibility as susceptible (S), intermediate (I), or resistant (R) presented in **Supplementary Table 3**.

Higher resistant patterns were shown for ampicillin, azithromycin, and nalidixic acid, with resistance rates exceeding 70%. Although cephalosporins and fluoroquinolones were shown to be intermediate in resistance, carbapenems (imipenem and meropenem) and chloramphenicol were among the most effective antibiotics (**Figure 1**).

**Figure 1.**
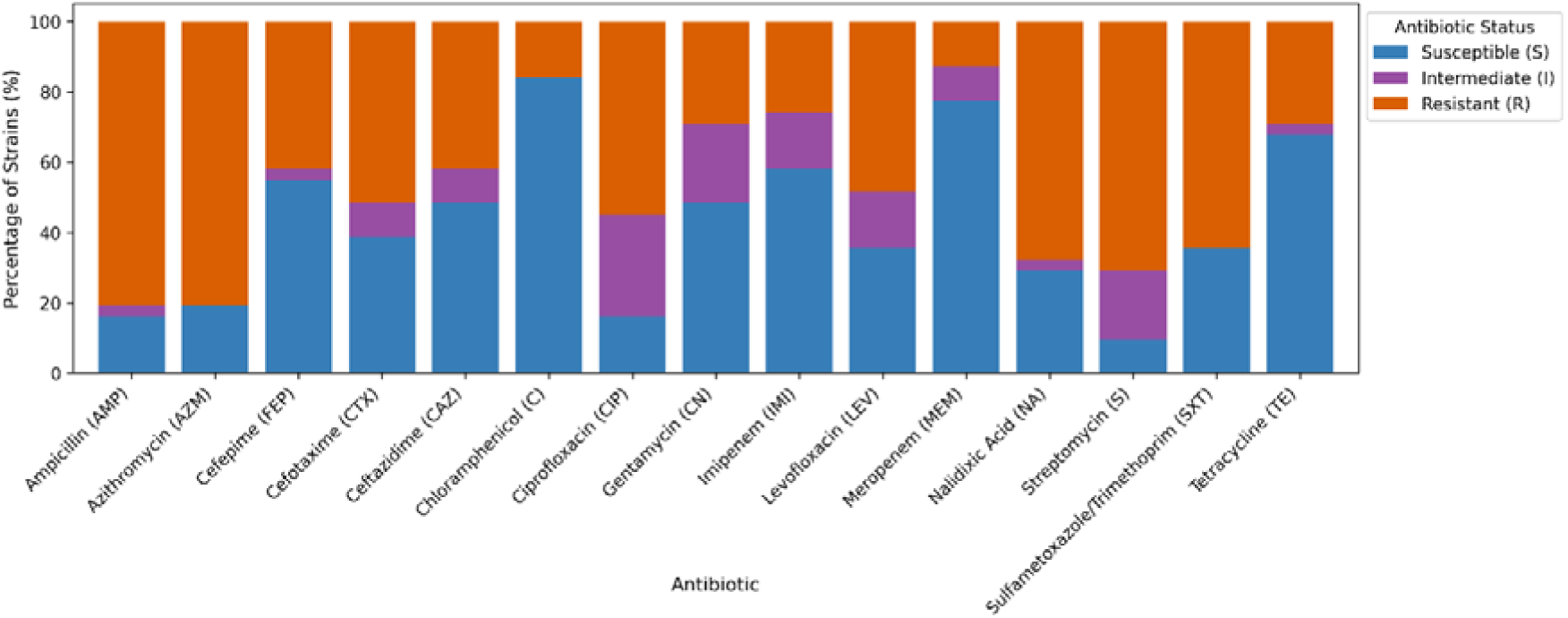
Antibiotic susceptibility of isolates against 15 antibiotics. Susceptible (S), Intermediate (I), Resistant (R) are indicated by blue, purple and orange respectively.

Strain-wise resistance distribution classified the isolates. Interestingly, NIB001, NIB002, and NIB006 were susceptible to nearly all tested antibiotics. Several strains (e.g., NIB009, NIB012, NIB015, NIB031, NIB037) were resistant to nearly all antibiotics tested which demonstrates the buildup of multi-drug resistance patterns within the patients **(Figure 2)**.

**Figure 2.**
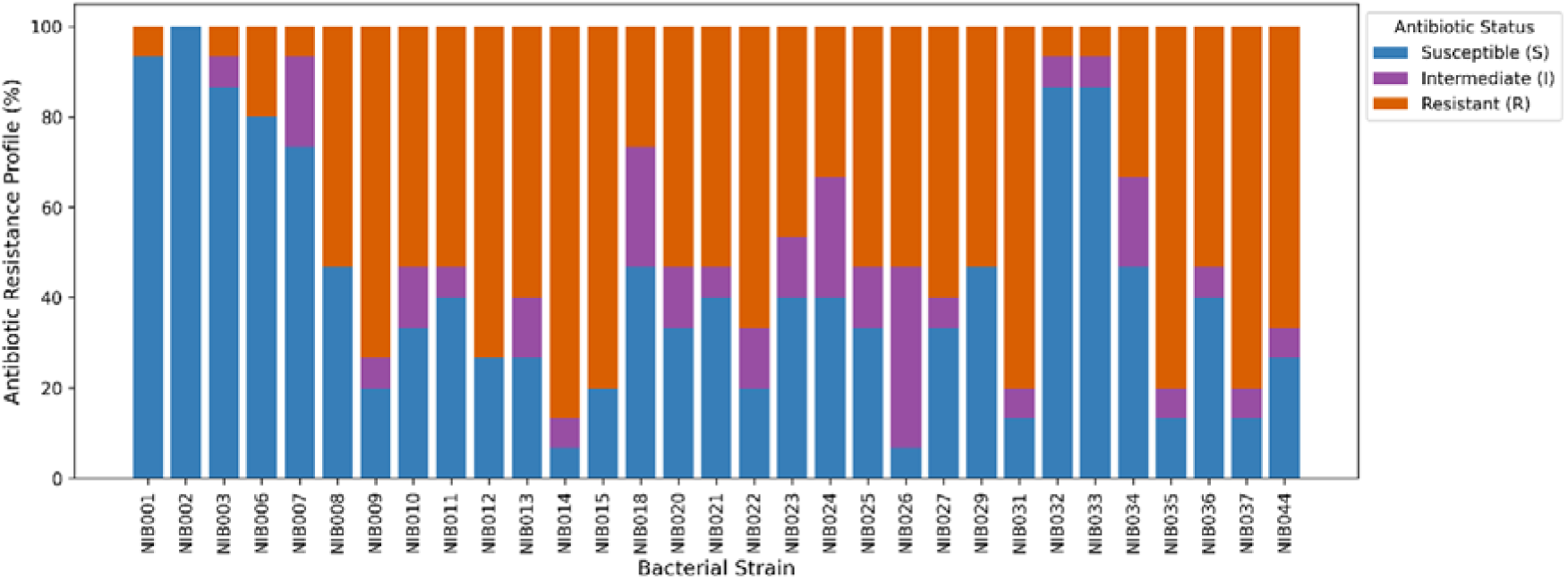
Strain-wise resistance profiles of bacterial isolates. Susceptible (S), Intermediate (I), Resistant (R) are indicated by blue, purple and orange respectively.

### Genomic features were revealed by assembly and annotation of the bacterial isolates

The assembly results indicated no presence of contaminating sequences from other organisms. The Unicycler Assembler was used to perform genome assembly of the ten bacterial isolates (*Citrobacter werkmanii, Enterobacter chuandaensis, Escherichia coli, Escherichia fergusonii, Enterobacter hormaechei, Klebsiella pneumoniae, Morganella morganii, Proteus mirabilis, Plesiomonas shigelloids* and *Providencia stuartii*) which provided high-quality draft genomes ranging from 3.6 Mb to 5.7 mb and contig counts between 8 to 388. The N50 values was an indicator of assembly quality, ranged from 15.9 Kb to 2.6 Mb. GC content was consistent ranging from 38.73% to 57.42%. The QUAST results (**Supplementary Data S4**) indicated no mismatches in these assembly. Assembled genome of bacterial isolates showed 3096 to 5303 coding DNA sequences (CDSs), 3 to 6 rRNAs, 45 to 94 tRNAs and 1 tmRNA in each isolate (**Supplementary Table S9**).

A complete graphical display of the distribution of the genome annotations of 10 bacterial isolates (*Citrobacter werkmanii, Enterobacter chuandaensis, Escherichia coli, Escherichia fergusonii, Enterobacter hormaechei, Klebsiella pneumoniae, Morganella morganii, Proteus mirabilis, Plesiomonas shigelloids, Providencia stuartii*) are provided in **Figure 3**.

**Figure 3:**
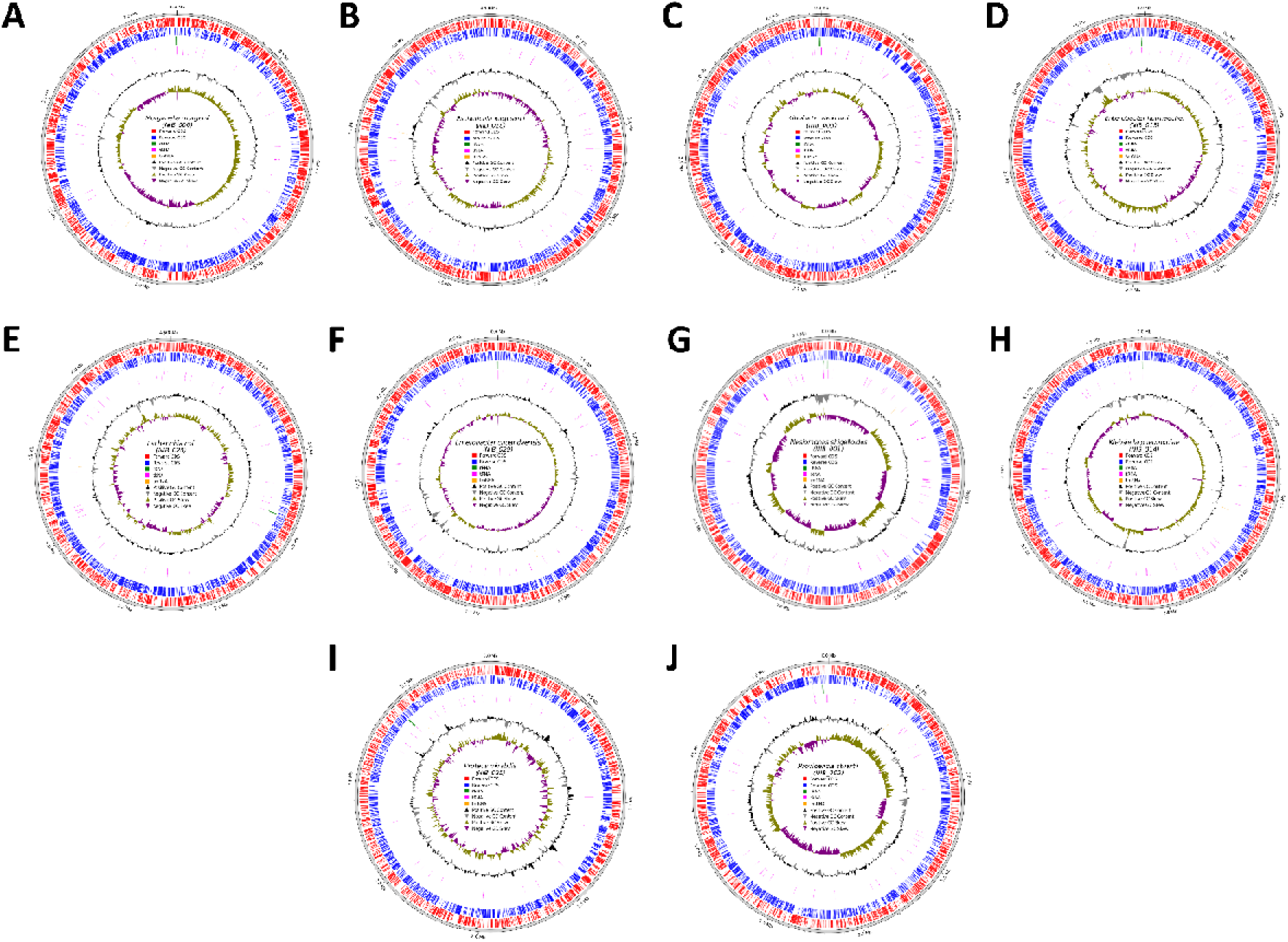
Complete genome annotation of *Morganella morganii* (**A**), *Escherichia fergusonii* (**B**), *Citrobacter werkmanii* (**C**), *Enterobacter hormaechei* (**D**), *Escherichia coli* (**E**), *Enterobacter chuandaensis* (**F**), *Plesiomonas shigelloids* (**G**), *Klebsiella pneumoniae* (**H**), *Proteus mirabilis* (**I**), *Providencia stuartii* (**J**).

### Pangenome analysis uncovers core genes and a diverse strain population

Out of the 100 bacterial genomes analysed (**Supplementary Table S10**), a total of 61,099 unique genes were identified. Among these, 50 genes were classified as core genes because they were present in at least 99% (n=100) of the strains (**Figure 4**). These core genes make up just 0.08% of the entire pangenome. Interestingly, no soft-core genes defined as genes present in 95-99% of the genomes were detected. The majority of the genes fell into the category of accessory genes, representing 99.92% of the pangenome. This included 5,781 shell genes and 55,268 cloud genes. Shell genes were found in 15-95% of the strains, while cloud genes were identified in fewer than 15% of the 100 strains.

**Figure 4.**
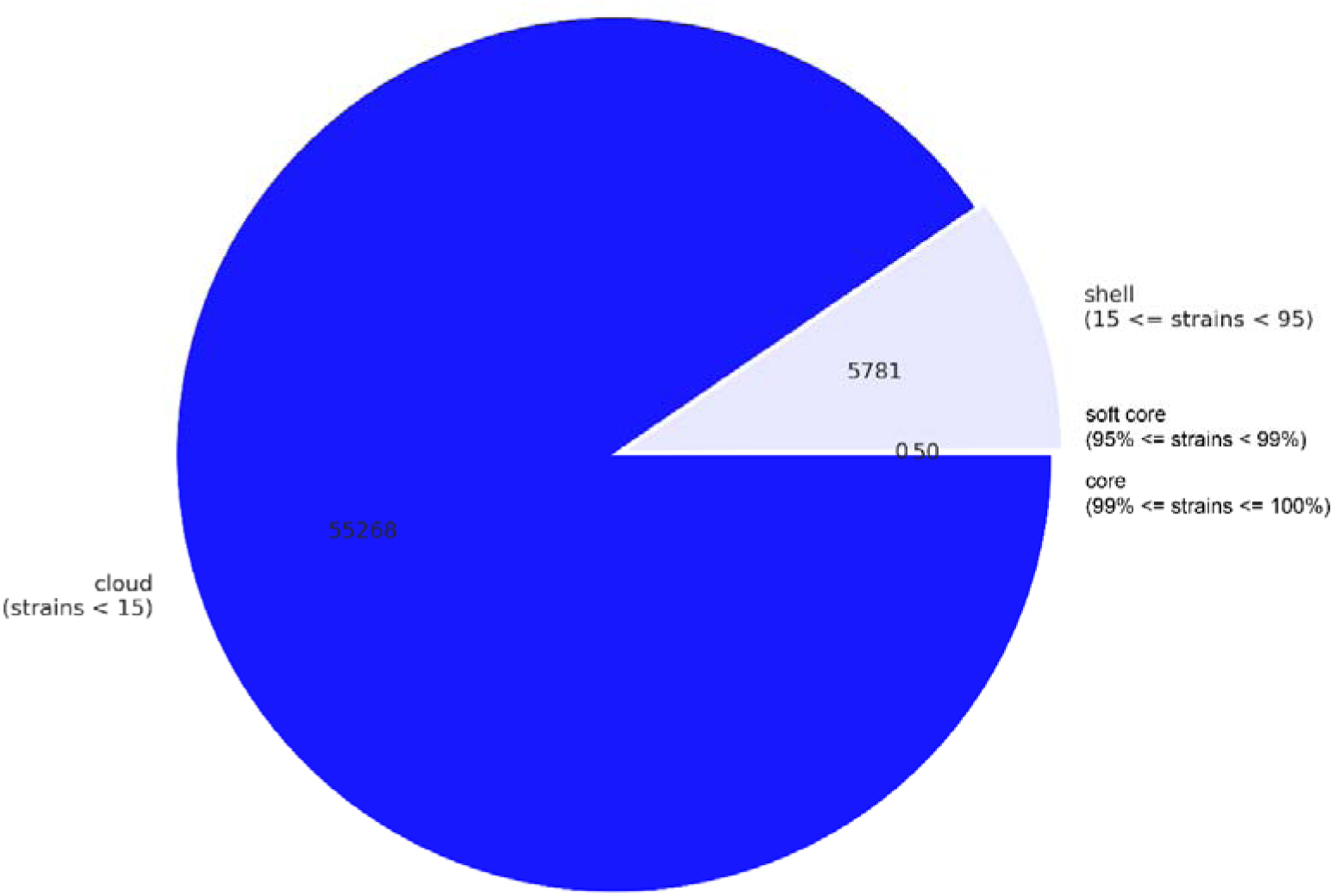
Distribution of Core, Soft-Core, Shell, and Cloud Genes within the Pangenome of the Analyzed Strains. The clusters consist of 50 core clusters, 55268 cloud clusters, and 5781 shell clusters of genes, with no soft-core gene clusters.

### Phylogenetic trees illustrated genetic variation among the genomes

The alignment of 50 core genes across 100 assembled and annotated genomes, along with the construction of a maximum-likelihood (ML) phylogenetic tree, revealed a diverse population structure. Most strains formed small clusters on distinct branches of the tree **(Figure 5)**. These clusters were categorized based on the presence or absence of the 50 core genes. The analysis identified a broad distribution of gene clusters throughout the phylogeny, with evidence of gene loss and gain observed in both closely related and distantly related lineages. Some lineages displayed significant gene loss, highlighting evolutionary divergence and genetic variability among the strains.

**Figure 5.**
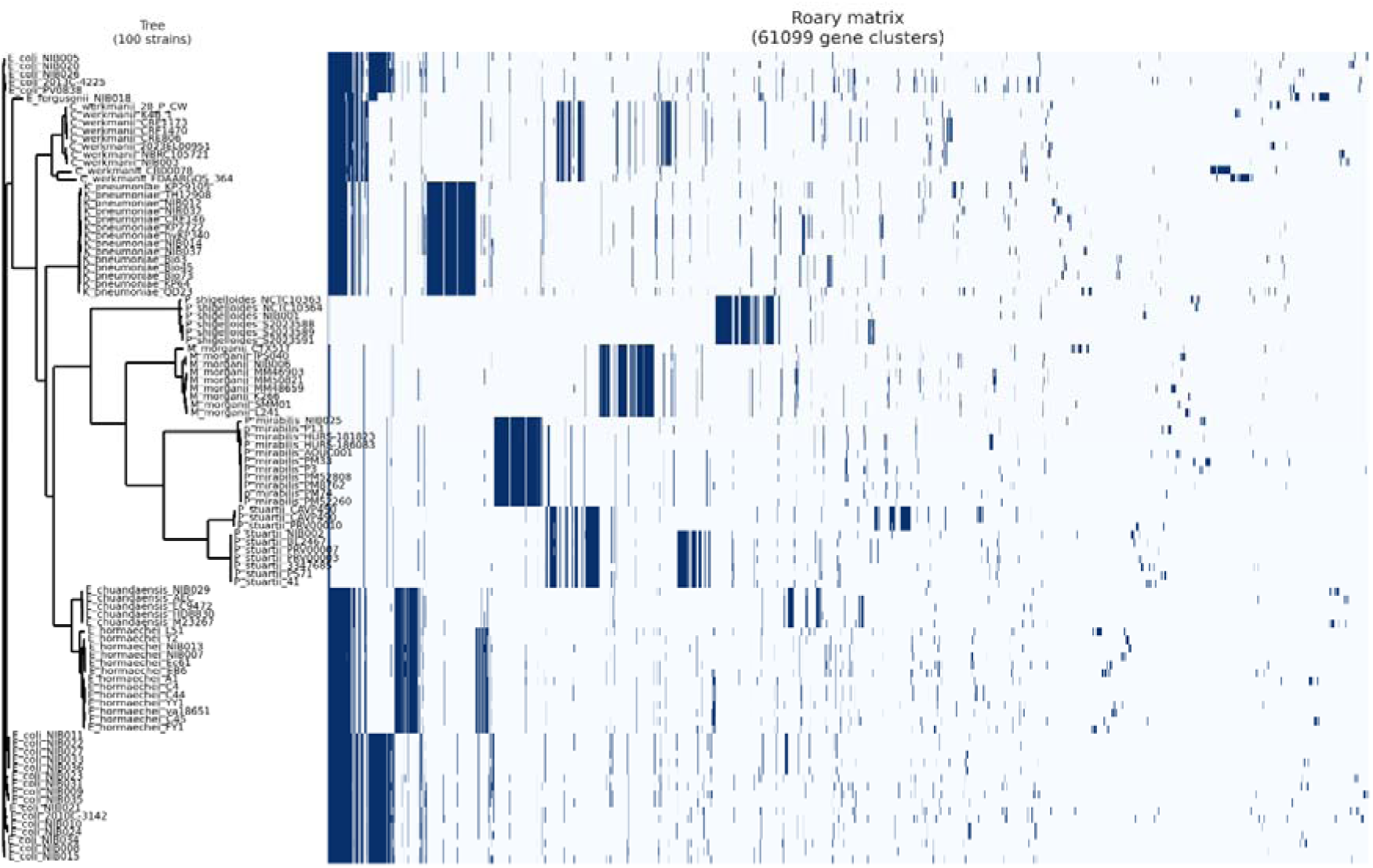
Maximum-likelihood phylogeny of 100 bacterial strains reconstructed using the 50 core-gene sequences identified in the pan-genomic analysis. A corresponding heat map was generated to visualize the distribution of 61,099 unique genes across the strains, with dark blue indicating the presence and white indicating the absence of each gene. The genes were arranged in columns, and their presence or absence was mapped onto the phylogenetic tree to highlight patterns of gene distribution among the strains.

### VFDB and CARD databases distinguished virulence and antimicrobial resistance gene of the bacterial isolates

By using ABricate with VFDB and CARD databases, we identified virulence and antimicrobial resistance (AMR) genes in 31 bacterial isolates from diarrheal patients. The most virulence and AMR genes were found in *Klebsiella pneumoniae*, followed by *Escherichia coli*, *Escherichia fergusonii*, *Citrobacter werkmanii*, *Enterobacter hormaechei*, and *Enterobacter cchandaensis*. *Plesiomonas shigelloides* had the fewest genes (2 virulence and 2 AMR), whereas *Proteus mirabilis, Morganella morganii,* and *Providencia stuartii* had no virulence genes and fewer AMR genes **(Table 3 & 4)**. The complete gene lists are presented in **Supplementary Table S5 and S6.**

**Table 3:**
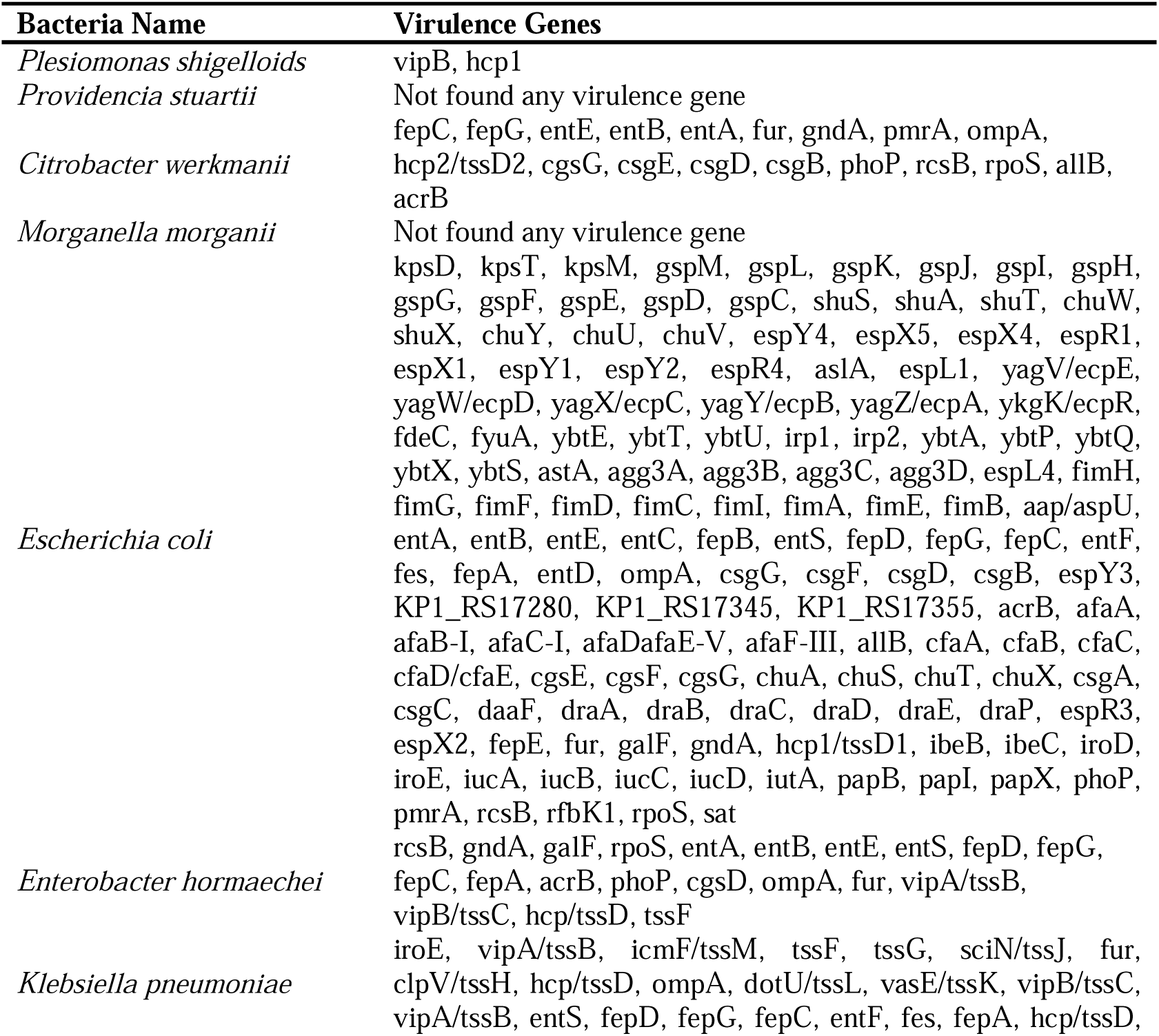
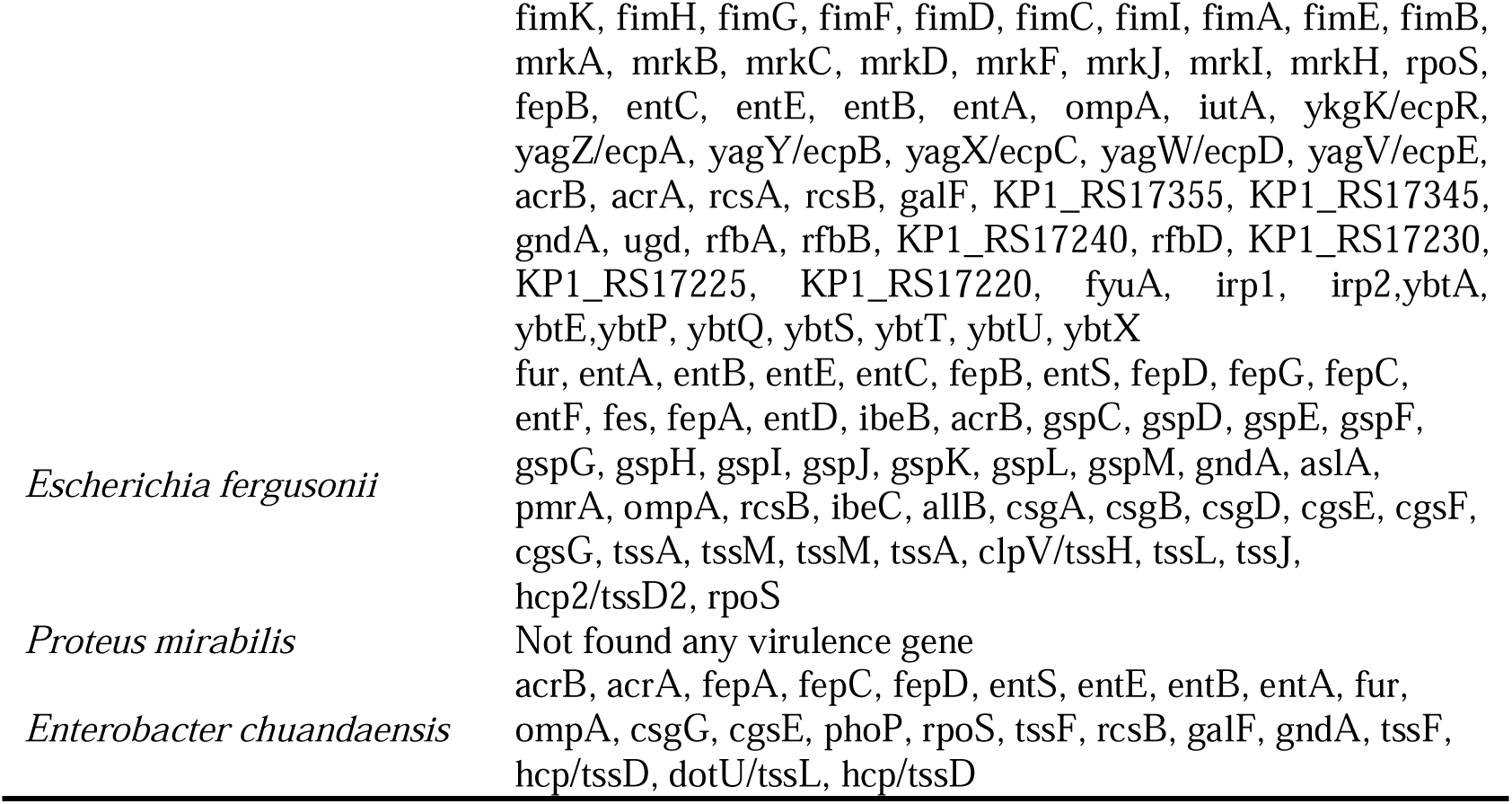
List of virulence genes of 10 unique bacterial isolates.

**Table 4:**
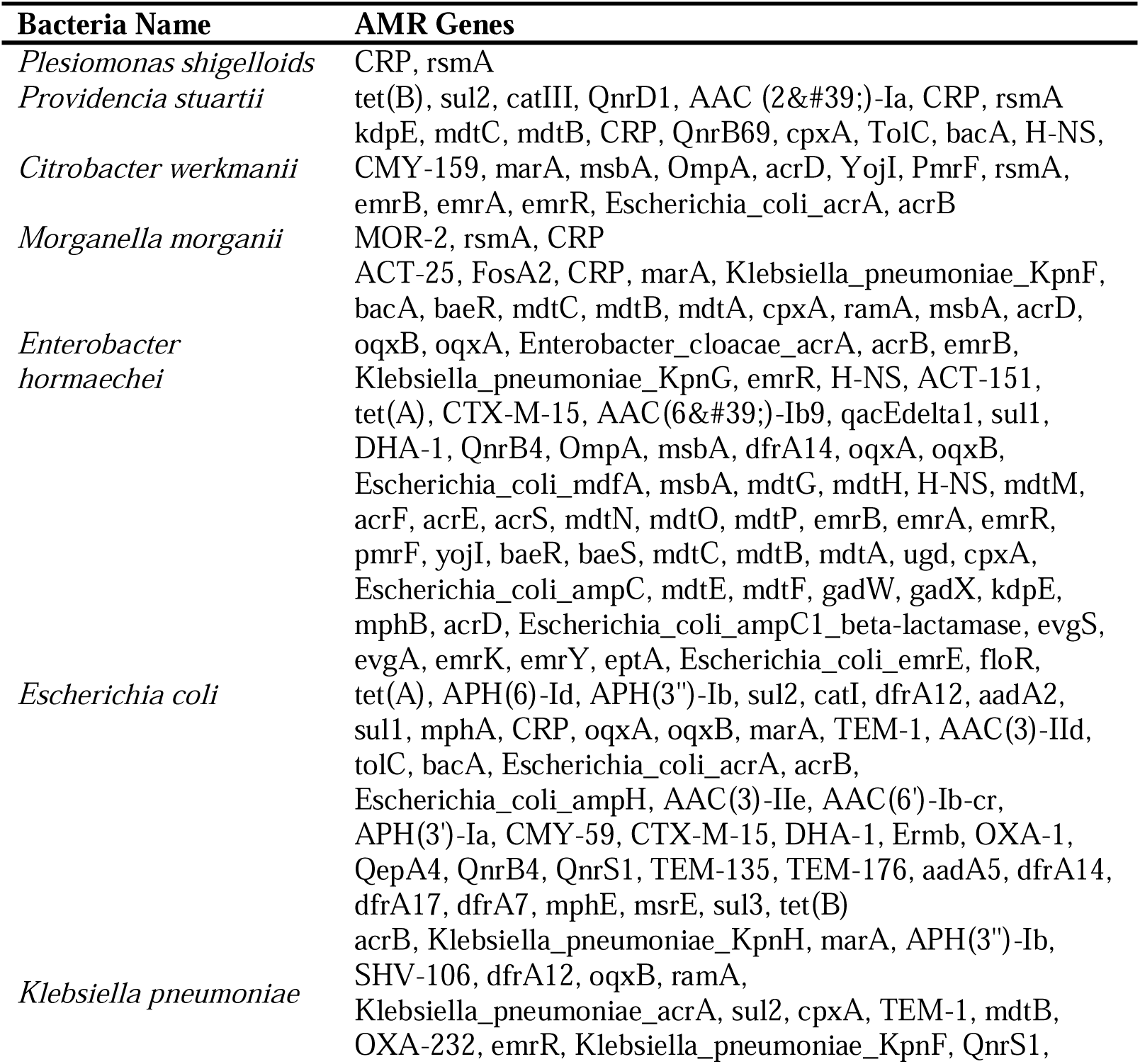
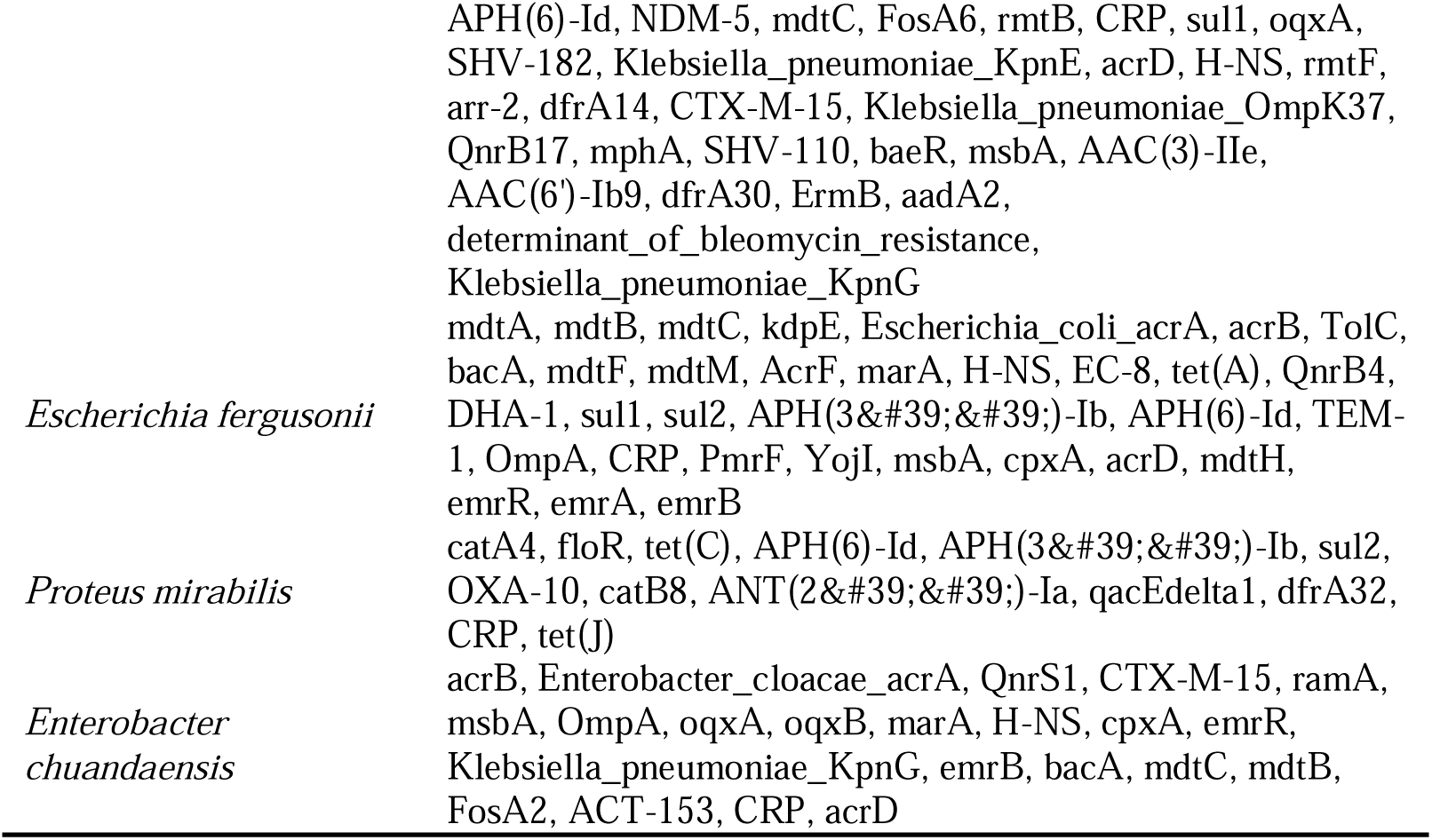
List of antimicrobial resistance genes of 10 unique bacterial isolates.

### Gene Ontology (GO) analysis of core genes defined functional characteristics

The Gene Ontology (GO) analysis **(Supplementary Table S7**) of 50 core genes by combining outputs from STRING, UniProt, KEGG pathway and RepSeq data demonstrated highly conserved functions which were common across the isolated bacterial species. The functions that were common included translation, gene expression, macromolecule biosynthetic process and other core functions.

The heatmap visualization (**Figure 6 and Supplementary Table S8**) further illustrated common functions of selected species by GO analysis.

**Figure 6.**
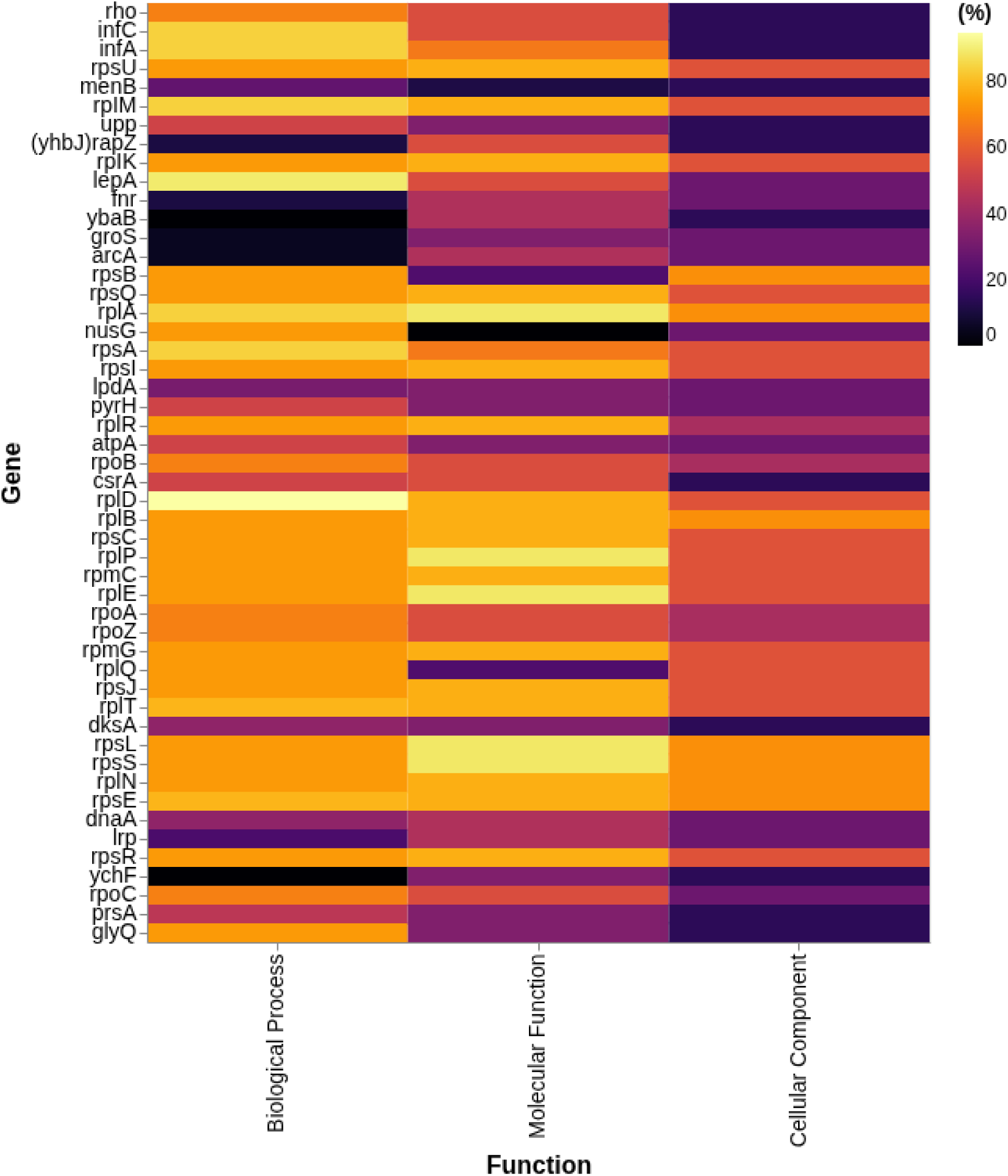
Heatmaps illustrating percentage of gene-functions common among the isolated species. Percentage of the gene-functions were plotted on X-axis and genes were plotted on Y-axis. Colour intensity changes from darker to lighter tone representing percentage of gene functions from low to high value (0% to 100%).

### Pathway Annotation Networks further visualized complex transcriptional dynamics of bacterial survival and pathogenicity

ClueGO pathway annotation analysis of core genes further elucidated the functional characteristics among the gene set and differentiated by percentages per group **(Figure 7)**. Highest percentage of genes (29.63%) shows the task of ribose phosphate biosynthetic process which is extremely crucial for nucleotide biosynthesis, bacterial replication and transcription. Upregulation of these genes further indicate the fitness of pathogenic strains during intestinal colonization. tRNA binding (16.05%) is another activity which is associated with these core genes indicates elevated virulent factors translation. Additionally, negative regulation of translation (11.11%) is another crucial activity conserving resources while ensuring production of essential factors. All of these activities associated with the core genes demonstrate the bacterial survival strategies in the host immune response while continued production of virulence factors. These activities are visualized by different clusters in **Figure 8**.

**Figure 7.**
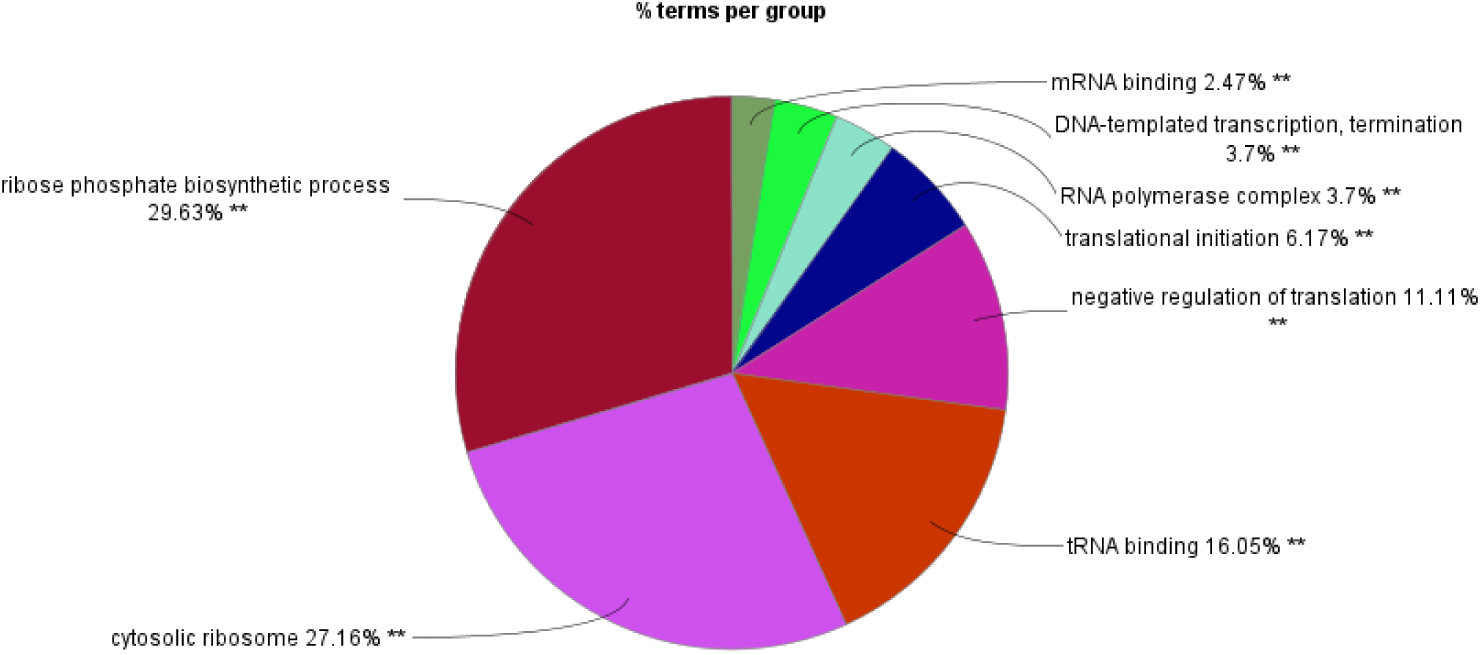
Pie chart shows percentage of core genes associated with each GO term related to Diarrhea. Highest percentage of genes (29.63%) have the activity of ribose phosphate biosynthetic process, tRNA binding (16.05%) and negative regulation of translation (11.11%) are followed after.

**Figure 8.**
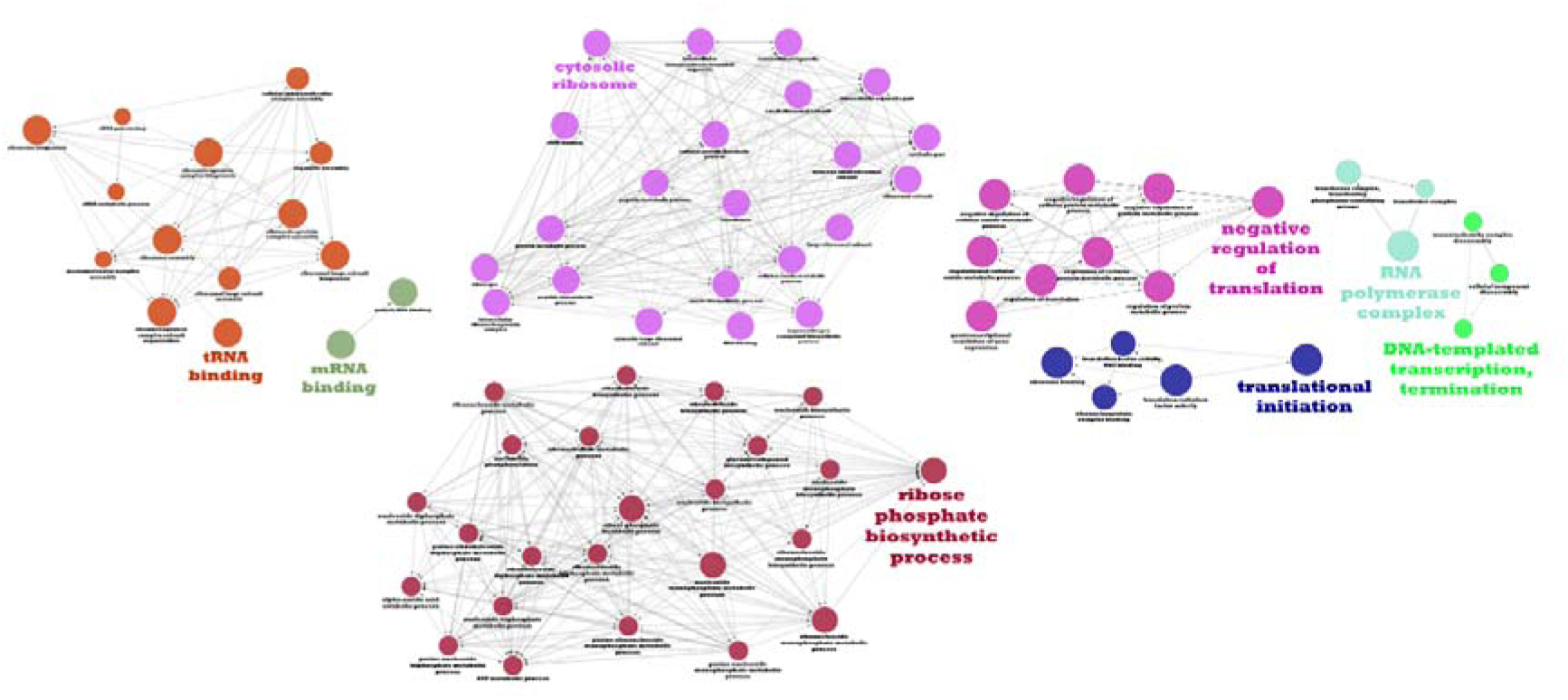
Pathway Annotation Network of core genes. It visualizes different crucial activities such as Ribosomal and Translational Activities, Metabolic Process, RNA Binding and Regulation, DNA Packaging and Replication Initiation in different clusters.

## Discussion

Herein, we describe the comprehensive pan-genome analysis following 31 gram-negative bacterial **(Table 1)** whole genome sequencing and provide the genomic characteristics, conferring insights **(Table 2 & Supplementary Table S3)** into bacterial survivability and pathogenicity. Moreover, we revealed the antibiotic resistance pattern of these sequenced bacteria **(Figure 1 & 2)**, which highlighted the growing concern for public health. Our pilot study provides a strong foundation for developing new therapeutics by exploring the whole genome, gaining pan-genomic insights, and furthering our understanding of antibiotic resistance patterns.

Previous studies on gram-negative bacteria of Bangladeshi isolates focused on their culture, molecular and biochemical identification (50,51), but our study extended the previous findings via the identification of critical genes for bacterial survival through the utilization of more advanced techniques.

Our study revealed significant genomic variability **(Figure 3 & Supplementary Table S2, S4)** in 10 unique gram-negative bacterial species isolated from diarrheal patients **(Supplementary Table S1)** in Bangladesh. Analyzing core and accessory genes reveals the genetic variety of bacterial strains and their dynamic genomic features **(Figure 4 & 5, Supplementary Table S9)**. A pan-genomic investigation of 100 WGS (31 clinical samples and 69 WGS from public databases) **(Supplementary Table S10)** revealed that nearly all strains conserved fifty (50) core genes. These identified core genes were all determined to be non-homologous which highlights the distinct and conserved roles that these genes play across different isolates. Among the core genes, ArcA, FNR, RapZ, csrA have activities in virulence and pathogenicity (52–55). The toxin screening did not uncover any genes in the core genome that encode toxins. This indicates that essential cellular and metabolic functions are maintained by core genes, while virulence determinants likely reside in shell and accessory regions. Furthermore, gene ontology (GO) analysis **(Figure 6 & Supplementary Table S7, S8)** provides substantial evidence of the functional roles of the core genes in basic processes like biosynthesis of macromolecules, translation, and cellular respiration **(Figure 7, 8)**, making them essential for bacteria’s survival and pathogenicity in the host’s gastrointestinal environment.

A high level of genomic plasticity is observed in accessory genes, which aids the notion that these pathogens have high adaptability, capable of obtaining and exchanging genetic materials associated with virulence and resistance. This kind of genomic plasticity may provide these pathogens with an edge in terms of their ability to survive, particularly when they are subjected to selective pressure from the immune cell responses of the host immune system and antimicrobial agents. Collectively, these findings provide valuable insights into the genomic basis of diarrheal disease and offer potential targets for future therapeutic interventions. Besides, we also provide insights on shell and accessory genes which revealed the presence of virulence-associated genes **(Table 3, Supplementary Table S5)** and antimicrobial resistance (AMR) genes **(Table 4, Supplementary Table S6).**

*Klebsiella pneumoniae* had the most virulence and AMR genes, followed by *E. coli, E. fergusonii, Citrobacter werkmanii, Enterobacter hormaechei* and *Enterobacter chuandaensis*. The identification of AMR genes suggests the potential for antibiotic resistance development, emphasizing the need for continuous surveillance. Moreover, virulence genes found within the shell and accessory genome may play a crucial role in host-pathogen interactions. Hence, the identification of accessory genes responsible for pathogen virulence suggests potential vaccine targets that can disrupt bacterial colony development and disease progression.

The presence of high number of AMR genes **(Supplementary Table S6)** in our isolates further led to explore on the antimicrobial resistance investigation to 15 antibiotics as AMR is alarming in Bangladesh. This growing concern is strengthened by the findings of our study, which revealed that particular isolates exhibited resistance to various medicines, including fluoroquinolones and beta-lactams. *Escherichia coli* demonstrated a high level of resistance to multiple antibiotics, including the critically essential carbapenems. These antibiotics are normally reserved for use as a last resort treatment for infections that do not respond to another antibiotic. This trend of resistance is consistent with the findings of global research (56) and it demonstrates how critical it is to combat antimicrobial resistance by enhancing antibiotic stewardship and discovering new therapeutics. Additionally, *Klebsiella pneumoniae* and *Enterobacter hormaechei* were found to be resistant to carbapenems. This is a reason for concern because it may indicate the presence of carbapenem-resistant *Enterobacteriaceae* (CRE). In hospitals and other medical facilities, microbes like these are recognized to be the source of infections. It is important to keep a close eye on how antimicrobial resistance is changing in these kinds of places so that immediate action can be performed while effective treatment options are accessible.

Hence, our study highlights the prospects of pan-genomic research in the development of targeted therapies for antibiotic-resistant infections. We can develop more effective treatments or diagnostic tools that can rapidly identify resistant strains by identifying genetic components associated with the immune system. This will allow for more effective and specific therapeutic targets.

## Limitations of the Study

Despite the significance of these findings from our study, there are some issues with our present study. Initially, we examined a limited number of samples, specifically 31 clinical samples. This may have induced bias, reduced the accuracy of our results. Further investigation is required using larger, more diverse sample populations from various global regions to validate our findings and assess the applicability of the identified core genes and resistance patterns across several contexts. This work focused solely on bacteria from children with diarrhoea; however, applying the same genomic analysis to adults might reveal the variations in bacterial populations and resistance profiles across different age cohorts.

## Conclusion

To summarize, this study provides a comprehensive genetic characterization of gram-negative bacteria associated with childhood diarrhoea in Bangladesh. We have identified core genes essential for bacterial survival and disease, as well as accessory genes that enhance the adaptability and resilience of these pathogens through whole-genome sequencing and pan-genome analysis. Increased antibiotic resistance has been identified in isolated bacteria, such as Escherichia coli and Klebsiella pneumoniae, highlighting the need for increased surveillance and the lookout for alternative treatment methods. The results of our study provide possible alternatives for future research, specifically focusing on the need to address the treatment gap of this disease by uncovering the underlying genetic mechanisms of diarrheal disease and antibiotic resistance. Continued genomic surveillance and targeted drug development are essential to address the increasing prevalence of diarrheal diseases in resource-limited countries like Bangladesh. Therefore, we believe that the information obtained through this research will play an important role in developing more effective prevention and treatment strategies in the future.

## Supporting information

S1. Demographic Data

S2. MALDI_TOF_result

S3. Antibiotic_Susceptibility_Data

S4. Assembly Quality

S5. Virulence Genes

S6. AMR_Genes_Details

S7. Gene_Ontology_Common_Functions

S8. Heatmap Data

S9. Genomic Assembly and Annotation Metrics

S10. List of Genomes for Pan-genome Analysis

## Declarations

### Ethics approval and consent to participate

We were authorized from the Institutional Review Board (IRB) of the National Institute of Biotechnology (NIB), Dhaka, Bangladesh (Approval No: NIB/IRB/2023/BID-04). This was done in accordance with ethical standards, and every individual who took part in this research was required to give their consent, either verbally or in writing.

### Consent for publication

Not applicable.

### Availability of data and materials

The original contributions presented in the study are publicly available through NCBI BioProjects. The strains are available as PRJNA1017991 (NIB001), PRJNA1017985 (NIB002), PRJNA1030290 (NIB003), PRJNA1187022 (NIB006), PRJNA1187023 (NIB007), PRJNA1308014 (NIB008), PRJNA1308304 (NIB009), PRJNA1308310 (NIB010), PRJNA1308487 (NIB011), PRJNA1308491 (NIB012), PRJNA1307279 (NIB013), PRJNA1190150 (NIB014), PRJNA1308000 (NIB015), PRJNA1308545 (NIB018), PRJNA1308774 (NIB020), PRJNA1308778 (NIB021), PRJNA1308580 (NIB022), PRJNA1308629 (NIB023), PRJNA1308794 (NIB024), PRJNA1308786 (NIB025), PRJNA1308796 (NIB026), PRJNA1308801 (NIB027), PRJNA1308802 (NIB029), PRJNA1309002 (NIB031), PRJNA1309013 (NIB032), PRJNA1309116 (NIB033), PRJNA1309124 (NIB034), PRJNA1309144 (NIB035), PRJNA1309147 (NIB036), PRJNA1309156 (NIB037), PRJNA1307376 (NIB044).

### Authors’ contributions

MUH, MAS, ZS, ANR conceptualized the project, carried out the laboratory experiments, analyzed sequence results and wrote the original draft. MNS and SSH analyzed the results, validated and visualized them with the assistance of MEH. ANR, MNS, SSH, MH and TP collected the clinical samples and processed them for microbiological analysis. MNS, ANA, AB, ZMC and IA reviewed and edited the original draft. MKH and PKS managed the resources. FMJ, MS and KCD supervised the project. All the authors approved the final draft.

## Acknowledgements

The authors are grateful to environmental, microbial, animal, and molecular biotechnology division of National Institute of Biotechnology for providing the lab access during this project.

