## Supplementary material for "Linking Genomic Landscape to Disease Mechanism: Core Genetic Factors Underlying Pathogenesis and Antimicrobial Resistance in Diarrheal Pathogens": S1. Demographic Data

| **Strain Number** | **Bacteria Name** | **Sex** | **Age (Months)** | **Location** | **NCBI BioProject** |
| --- | --- | --- | --- | --- | --- |
| NIB001 | *Plesiomonas shigelloids* | Female | 96 | Dhaka Shishu Hospital, Dhaka | PRJNA1017991 |
| NIB002 | *Providencia stuartii* | Male | 30 | Dhaka Shishu Hospital, Dhaka | PRJNA1017985 |
| NIB003 | *Citrobacter werkmanii* | Male | 6 | Dhaka Shishu Hospital, Dhaka | PRJNA1030290 |
| NIB006 | *Morganella morganii* | Male | 54 | Dhaka Shishu Hospital, Dhaka | PRJNA1187022 |
| NIB007 | *Enterobacter hormaechei* | Female | 96 | Dhaka Shishu Hospital, Dhaka | PRJNA1187023 |
| NIB008 | *Escherichia coli* | Female | 18 | Lakshmipur Sadar Hospital, Lakshmipur | PRJNA1308014 |
| NIB009 | *Escherichia coli* | Male | 13 | Lakshmipur Sadar Hospital, Lakshmipur | PRJNA1308304 |
| NIB010 | *Escherichia coli* | Female | 24 | Lakshmipur Sadar Hospital, Lakshmipur | PRJNA1308310 |
| NIB011 | *Escherichia coli* | Male | 8 | Dhaka Shishu Hospital & Institute | PRJNA1308487 |
| NIB012 | *Klebsiella pneumoniae* | Male | 24 | Lakshmipur Sadar Hospital, Lakshmipur | PRJNA1308491 |
| NIB013 | *Enterobacter hormaechei* | Male | 8 | Dhaka Shishu Hospital & Institute | PRJNA1307279 |
| NIB014 | *Klebsiella pneumoniae* | Male | 8 | Dhaka Shishu Hospital & Institute | PRJNA1190150 |
| NIB015 | *Escherichia coli* | Male | 4 | Dhaka Shishu Hospital & Institute | PRJNA1308000 |
| NIB018 | *Escherichia fergusonii* | Female | 70 | Lakshmipur Sadar Hospital, Lakshmipur | PRJNA1308545 |
| NIB020 | *Escherichia coli* | Female | 18 | Lakshmipur Sadar Hospital, Lakshmipur | PRJNA1308774 |
| NIB021 | *Escherichia coli* | Female | 8 | Dhaka Shishu Hospital, Dhaka | PRJNA1308778 |
| NIB022 | *Escherichia coli* | Male | 22 | Dhaka Shishu Hospital, Dhaka | PRJNA1308580 |
| NIB023 | *Escherichia coli* | Male | 8 | Dhaka Shishu Hospital, Dhaka | PRJNA1308629 |
| NIB024 | *Escherichia coli* | Female | 24 | Lakshmipur Sadar Hospital, Lakshmipur | PRJNA1308794 |
| NIB025 | *Proteus mirabilis* | Female | 10 | Dhaka Shishu Hospital, Dhaka | PRJNA1308786 |
| NIB026 | *Escherichia coli* | Male | 24 | Lakshmipur Sadar Hospital, Lakshmipur | PRJNA1308796 |
| NIB027 | *Escherichia coli* | Female | 12 | Dhaka Shishu Hospital, Dhaka | PRJNA1308801 |
| NIB029 | *Enterobacter chuandaensis* | Male | 24 | Lakshmipur Sadar Hospital, Lakshmipur | PRJNA1308802 |
| NIB031 | *Escherichia coli* | Female | 8 | Lakshmipur Sadar Hospital, Lakshmipur | PRJNA1309002 |
| NIB032 | *Klebsiella pneumoniae* | Male | 22 | Dhaka Shishu Hospital, Dhaka | PRJNA1309013 |
| NIB033 | *Escherichia coli* | Female | 18 | Lakshmipur Sadar Hospital, Lakshmipur | PRJNA1309116 |
| NIB034 | *Escherichia coli* | Male | 7 | Dhaka Shishu Hospital, Dhaka | PRJNA1309124 |
| NIB035 | *Escherichia coli* | Female | 6 | Lakshmipur Sadar Hospital, Lakshmipur | PRJNA1309144 |
| NIB036 | *Escherichia coli* | Female | 12 | Dhaka Shishu Hospital, Dhaka | PRJNA1309147 |
| NIB037 | *Klebsiella pneumoniae* | Male | 9 | Dhaka Shishu Hospital, Dhaka | PRJNA1309156 |
| NIB044 | *Escherichia coli* | Male | 14 | Lakshmipur Sadar Hospital, Lakshmipur | PRJNA1307376 |
