## Supplementary material for "Linking Genomic Landscape to Disease Mechanism: Core Genetic Factors Underlying Pathogenesis and Antimicrobial Resistance in Diarrheal Pathogens": S2. MALDI_TOF_result

| **Strain Number** | | **Rank (Quality)** | **Top Match Organism** | | **Score Value** |
| --- | --- | --- | --- | --- | --- |
| NIB001 | | (+++) | *Plesiomonas shigelloids* | | 2.13 |
| NIB002 | | (+) | *Providencia stuartii* | | 1.91 |
| NIB003 | | (+++) | *Citrobacter werkmanii* | | 2.13 |
| NIB006 | | (+) | *Morganella morganii* | | 1.99 |
| NIB007 | | (+) | *Enterobacter hormaechei* | | 1.9 |
| NIB008 | | (+++) | *Escherichia coli* | | 2.03 |
| NIB009 | | (+++) | *Escherichia coli* | | 2.01 |
| NIB010 | | (+++) | *Escherichia coli* | | 2.11 |
| NIB011 | | (+) | *Escherichia coli* | | 1.83 |
| NIB012 | | (+++) | *Klebsiella pneumoniae* | | 2.33 |
| NIB013 | | (+++) | *Enterobacter hormaechei* | | 2.33 |
| NIB014 | | (+++) | *Klebsiella pneumoniae* | | 2.44 |
| NIB015 | | (+++) | *Escherichia coli* | | 2.22 |
| NIB018 | | (+++) | *Escherichia fergusonii* | | 2.21 |
| NIB020 | | (+++) | *Escherichia coli* | | 2.12 |
| NIB021 | | (+) | *Escherichia coli* | | 1.97 |
| NIB022 | | (+++) | *Escherichia coli* | | 2.18 |
| NIB023 | | (+++) | *Escherichia coli* | | 2.32 |
| NIB024 | | (+++) | *Escherichia coli* | | 2.27 |
| NIB025 | | (+++) | *Proteus mirabilis* | | 2.05 |
| NIB026 | | (+++) | *Escherichia coli* | | 2.16 |
| NIB027 | | (+++) | *Escherichia coli* | | 2.13 |
| NIB029 | | (+++) | *Enterobacter chuandaensis* | | 2.2 |
| NIB031 | | (+++) | *Escherichia coli* | | 2.23 |
| NIB032 | | (+) | *Klebsiella pneumoniae* | | 1.76 |
| NIB033 | | (+++) | *Escherichia coli* | | 2.01 |
| NIB034 | | (+++) | *Escherichia coli* | | 2.23 |
| NIB035 | | (+++) | *Escherichia coli* | | 2.11 |
| NIB036 | | (+++) | *Escherichia coli* | | 2.3 |
| NIB037 | | (+++) | *Klebsiella pneumoniae* | | 2.26 |
| NIB044 | | (+++) | *Escherichia coli* | | 2.21 |
| **Note:** |  |  |  |  |  |
| Range 2.00-3.00 = High-confidence identification (+++) | | | |  |  |
| Range 1.70-1.99 = Low-confidence identification (+) | | | |  |  |
| Range 0.00-1.69 = No organism identification possible (-) | | | |  |  |
