## Supplementary material for "Linking Genomic Landscape to Disease Mechanism: Core Genetic Factors Underlying Pathogenesis and Antimicrobial Resistance in Diarrheal Pathogens": S3. Antibiotic_Susceptibility_Data

| **Bacteria Name** | **Strain Number** | **Antibiotic Name** | | | | | | | | | | | | | | |
| --- | --- | --- | --- | --- | --- | --- | --- | --- | --- | --- | --- | --- | --- | --- | --- | --- |
|  |  | **Ampicillin (AMP)** | **Cefotaxime (CTX)** | **Ceftazidime (CAZ)** | **Gentamycin (CN)** | **Streptomycin (S)** | **Tetracycline (TE)** | **Chloramphenicol (C)** | **Ciprofloxacin (CIP)** | **Levofloxacin (LEV)** | **Nalidixic Acid (NA)** | **Sulfametoxazole/Trimethoprim (SXT)** | **Meropenem (MEM)** | **Imipenem (IMI)** | **Azithromycin (AZM)** | **Cefepime (FEP)** |
| *Plesiomonas shigelloides* | NIB001 | S | S | S | S | R | S | S | S | S | S | S | S | S | S | S |
| *Providencia stuartii* | NIB002 | S | S | S | S | S | S | S | S | S | S | S | S | S | S | S |
| *Citrobacter werkmanii* | NIB003 | S | S | S | S | R | S | S | I | S | S | S | S | S | S | S |
| *Morganella morganii* | NIB006 | S | S | S | S | S | R | R | S | S | S | S | S | S | R | S |
| *Enterobacter hormaechei* | NIB007 | I | S | S | S | I | S | S | S | S | I | S | S | S | R | S |
| *Escherichia coli* | NIB008 | R | S | S | S | R | R | S | R | R | R | R | S | S | R | S |
| *Escherichia coli* | NIB009 | R | R | R | I | R | R | S | R | R | R | R | S | S | R | R |
| *Escherichia coli* | NIB010 | R | I | S | I | R | S | S | R | R | R | R | S | R | R | S |
| *Escherichia coli* | NIB011 | R | R | R | S | R | S | S | I | S | R | R | S | S | R | R |
| *Klebsiella pneumoniae* | NIB012 | R | R | R | R | R | S | S | R | R | R | R | S | S | R | R |
| *Enterobacter hormaechei* | NIB013 | R | S | S | R | R | R | R | I | I | R | R | S | R | R | S |
| *Klebsiella pneumoniae* | NIB014 | R | R | R | R | I | S | R | R | R | R | R | R | R | R | R |
| *Escherichia coli* | NIB015 | R | R | R | R | R | S | S | R | R | R | R | S | R | R | R |
| *Escherichia fergusonii* | NIB018 | R | S | S | I | R | R | S | I | I | S | S | S | I | R | S |
| *Escherichia coli* | NIB020 | R | S | S | I | R | R | S | R | R | R | R | S | I | R | S |
| *Escherichia coli* | NIB021 | R | S | S | R | R | I | R | R | R | R | R | S | S | S | S |
| *Escherichia coli* | NIB022 | R | R | R | R | R | R | S | I | I | S | R | R | R | R | S |
| *Escherichia coli* | NIB023 | R | R | I | S | R | S | S | R | R | R | S | I | S | R | S |
| *Escherichia coli* | NIB024 | R | I | I | S | I | S | S | R | S | R | R | S | I | R | S |
| *Proteus mirabilis* | NIB025 | R | S | S | R | R | R | R | I | I | R | R | S | S | R | S |
| *Escherichia coli* | NIB026 | R | I | I | I | R | R | S | R | R | R | R | I | I | R | I |
| *Escherichia coli* | NIB027 | R | R | R | S | R | S | S | R | I | R | R | S | S | R | R |
| *Enterobacter chuandaensis* | NIB029 | R | R | S | S | R | S | S | R | R | R | S | S | S | R | R |
| *Escherichia coli* | NIB031 | R | R | R | I | R | S | S | R | R | R | R | R | R | R | R |
| *Klebsiella pneumoniae* | NIB032 | R | S | S | S | I | S | S | S | S | S | S | S | S | S | S |
| *Escherichia coli* | NIB033 | S | R | S | S | S | S | S | I | S | S | S | S | S | S | S |
| *Escherichia coli* | NIB034 | R | R | R | I | I | S | S | I | S | S | S | S | S | R | R |
| *Escherichia coli* | NIB035 | R | R | R | R | R | S | S | R | R | R | R | I | R | R | R |
| *Escherichia coli* | NIB036 | R | R | R | S | R | S | S | I | S | R | R | S | S | R | R |
| *Klebsiella pneumoniae* | NIB037 | R | R | R | R | I | S | S | R | R | R | R | R | R | R | R |
| *Escherichia coli* | NIB044 | R | R | R | S | R | S | S | R | R | R | R | S | I | R | R |
