## Supplementary material for "Linking Genomic Landscape to Disease Mechanism: Core Genetic Factors Underlying Pathogenesis and Antimicrobial Resistance in Diarrheal Pathogens": S4. Assembly Quality

**B1: Assembly quality of the *Plesiomonas shigelloides***

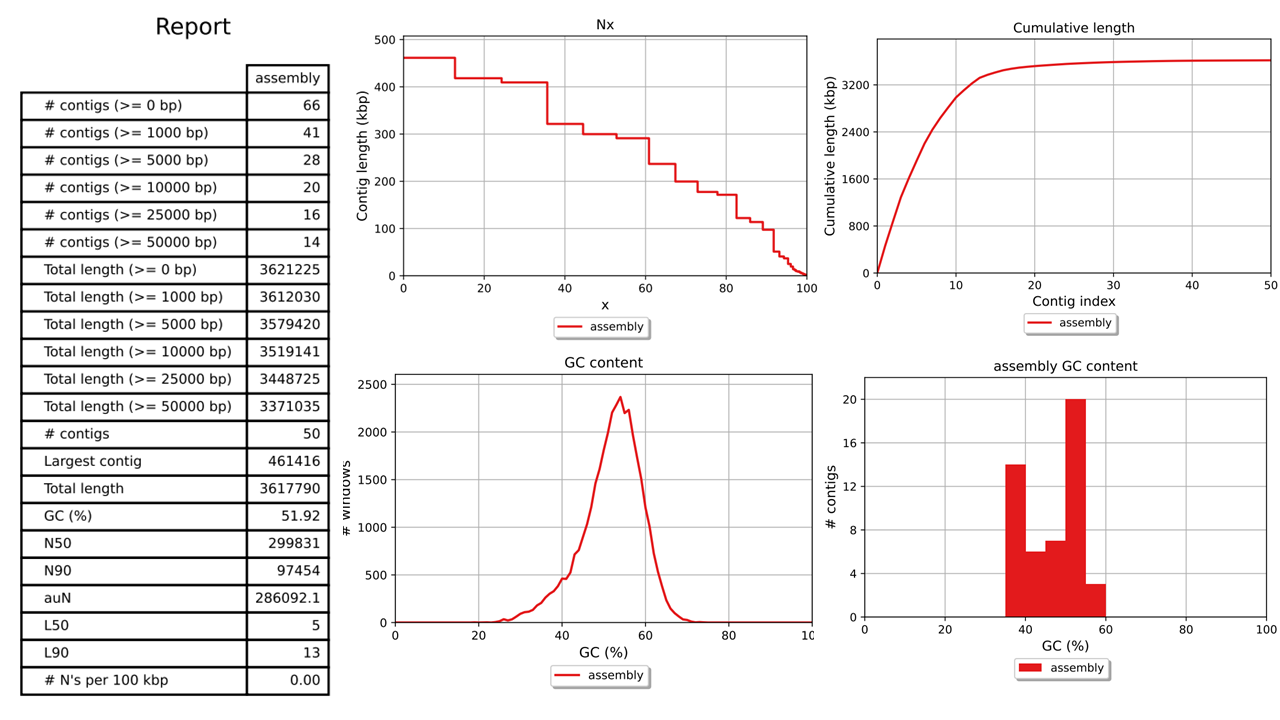

**B2: Assembly quality of the *Escherichia fergusonii***

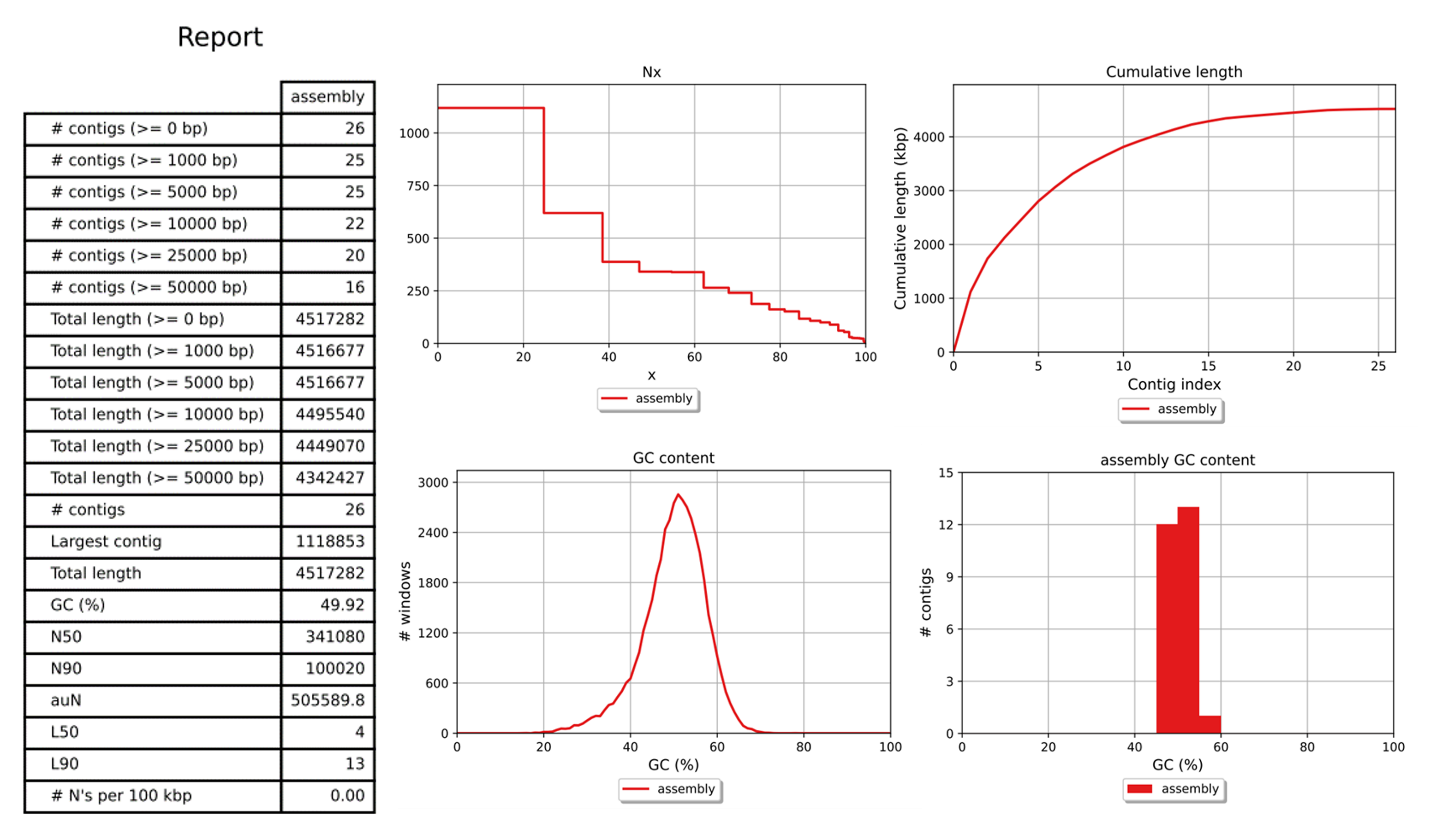

**B3: Assembly quality of the *Enterobacter chuandaensis***

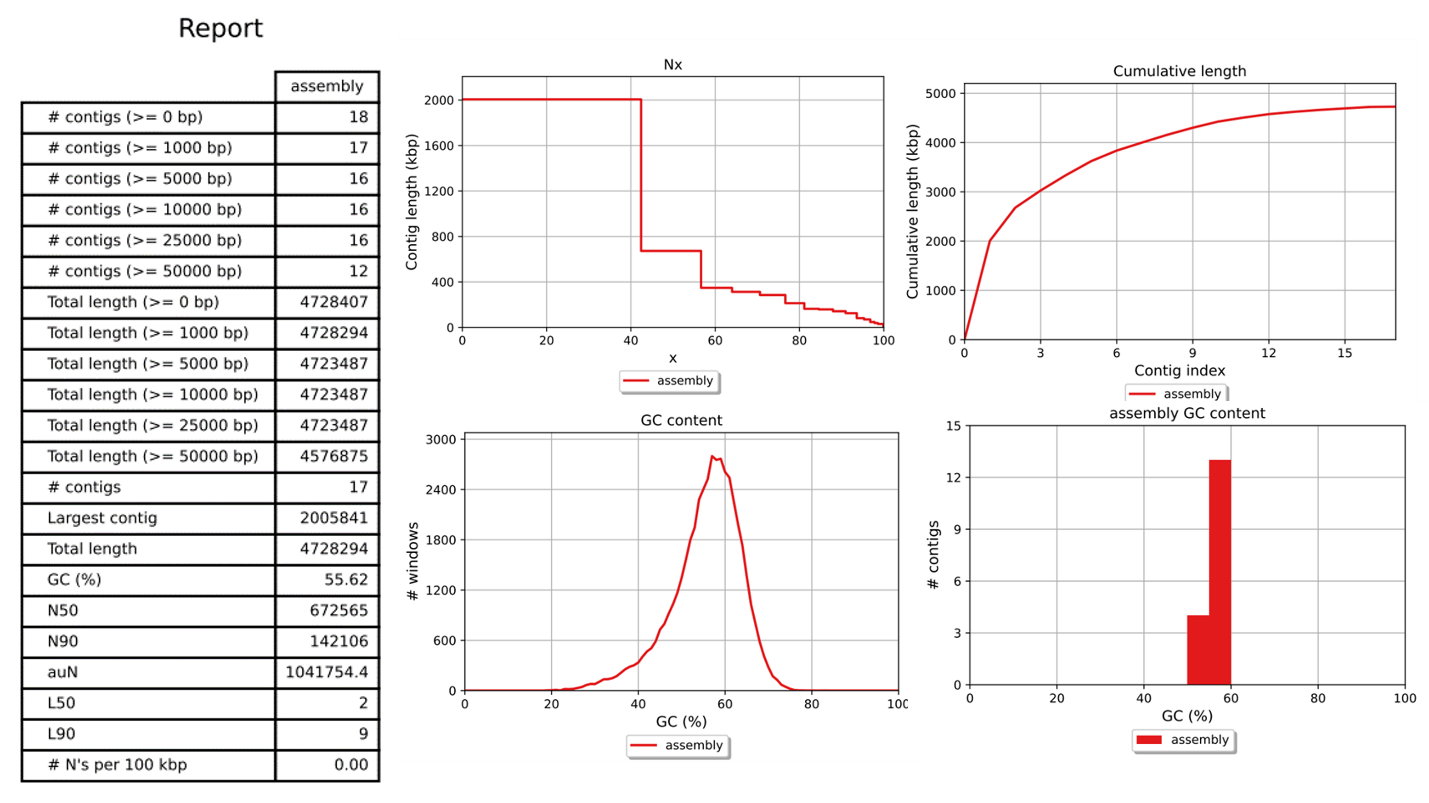

**B4: Assembly quality of the *Escherichia coli***

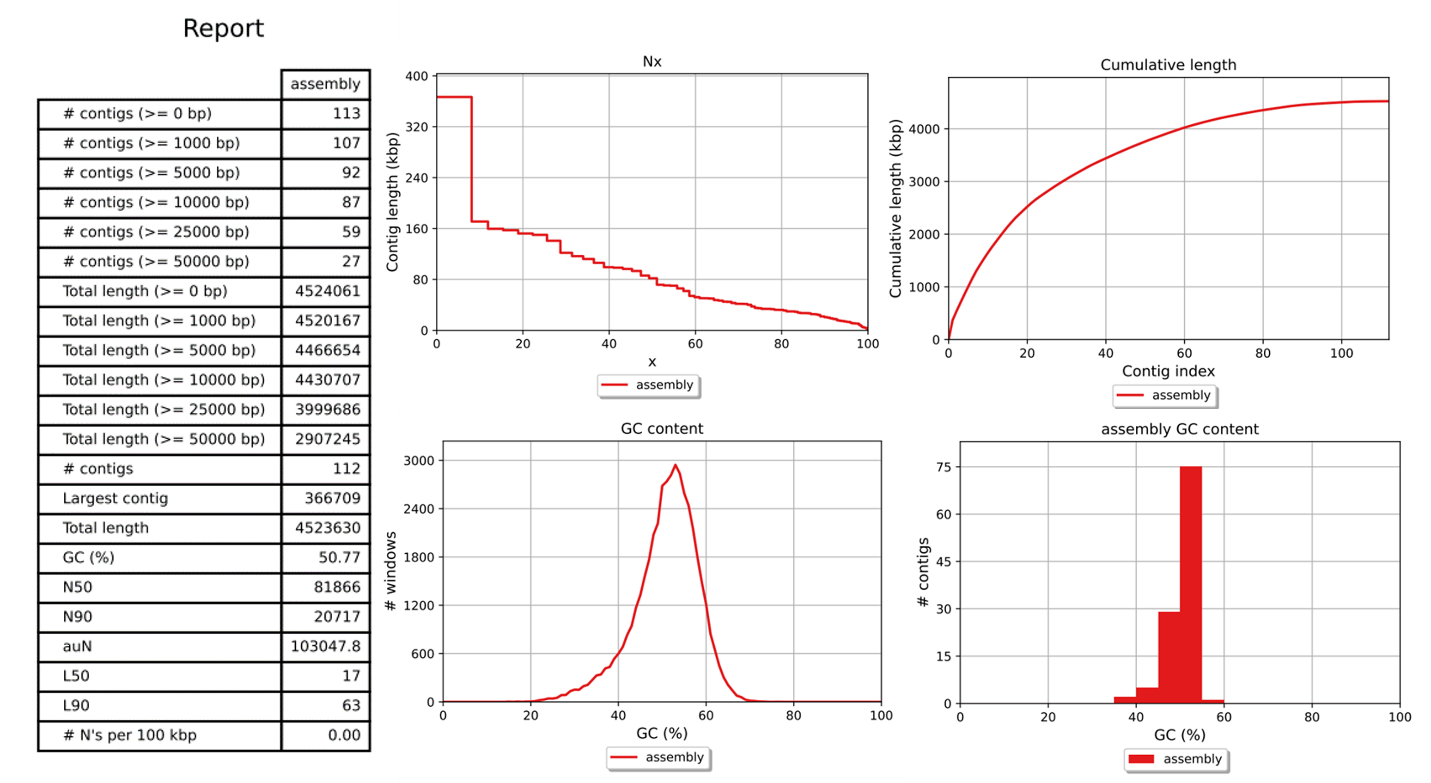

**B5: Assembly quality of the *Klebsiella pneumoniae***

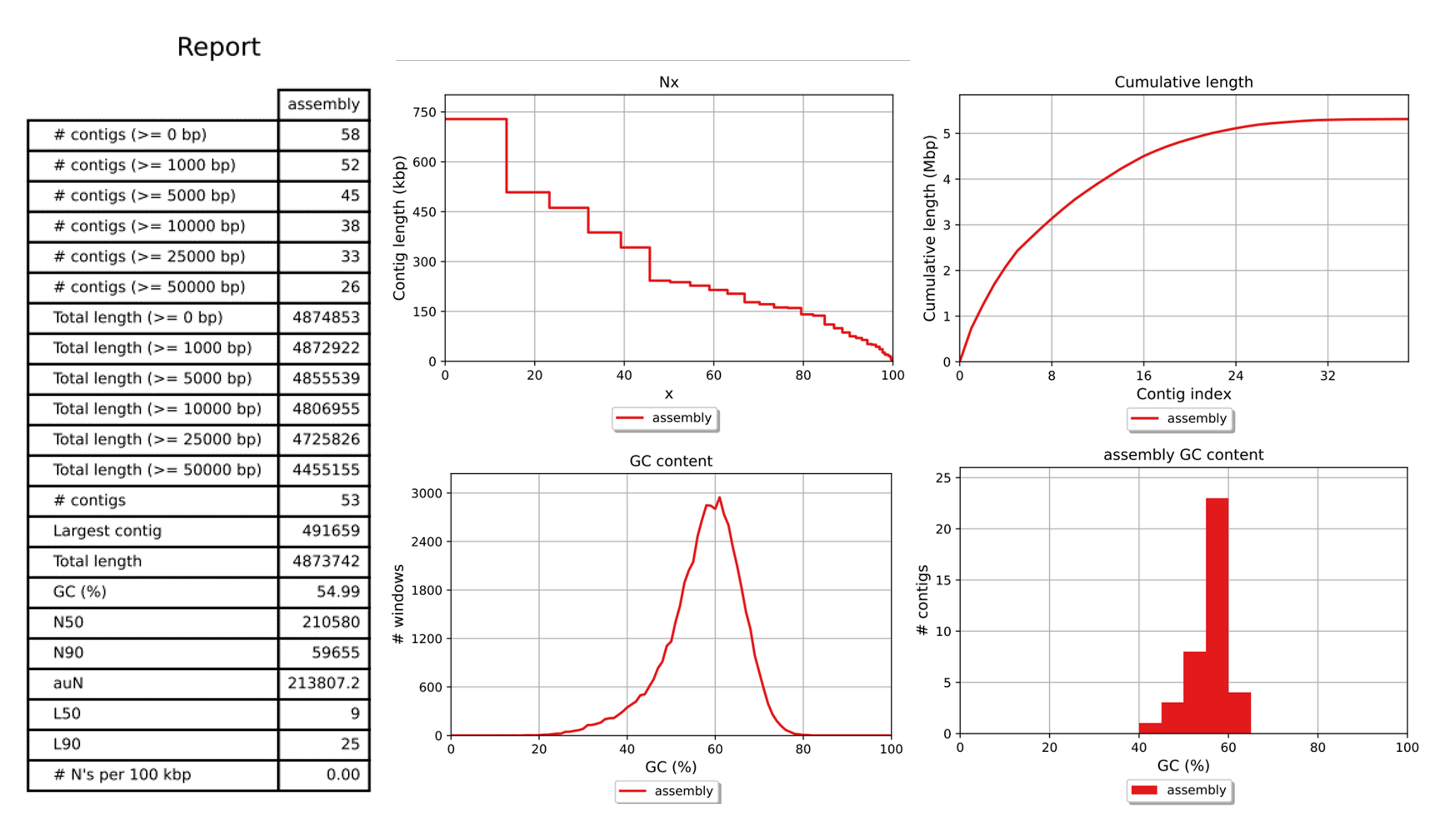

**B6: Assembly quality of the *Morganella morganii***

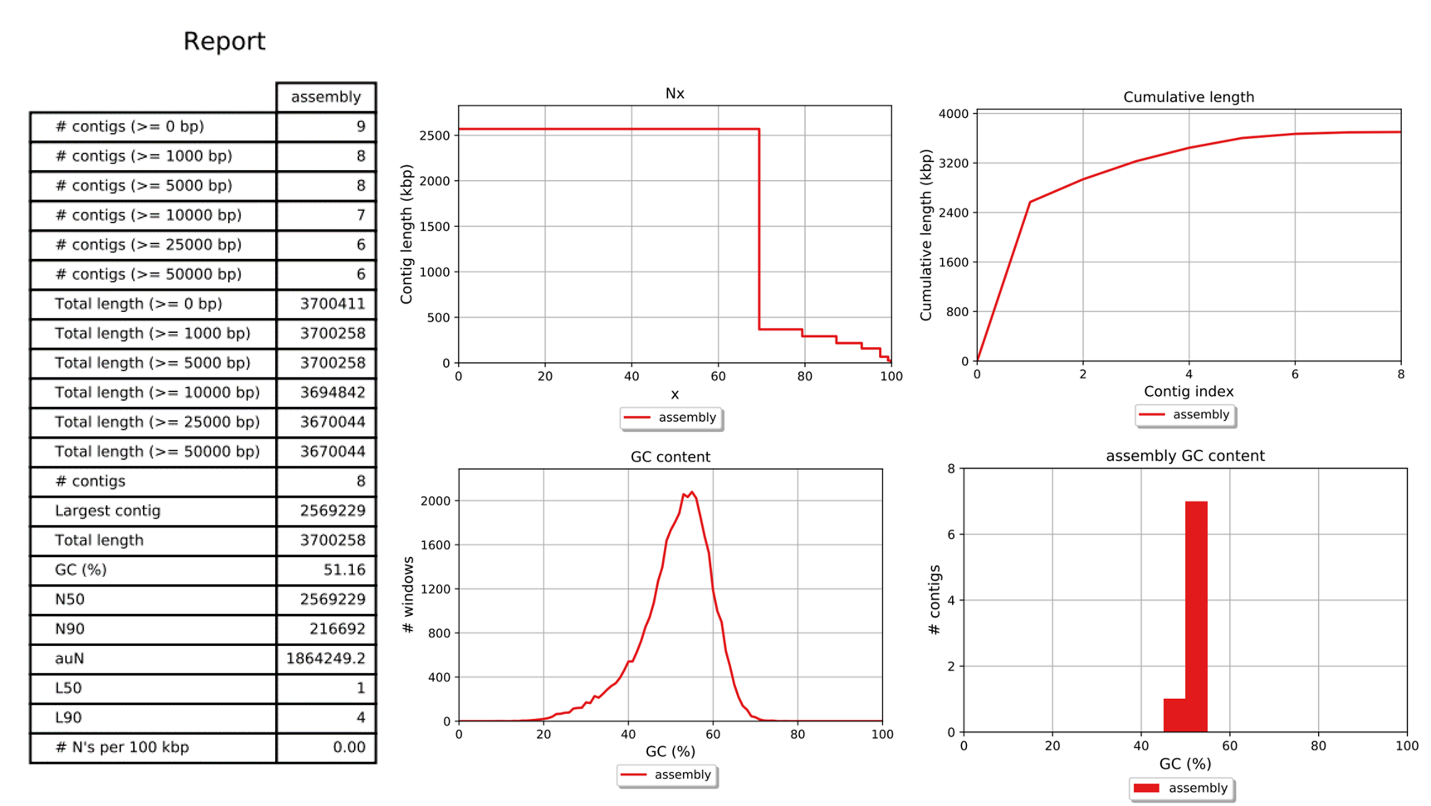

**B7: Assembly quality of the *Enterobacter hormaechei***

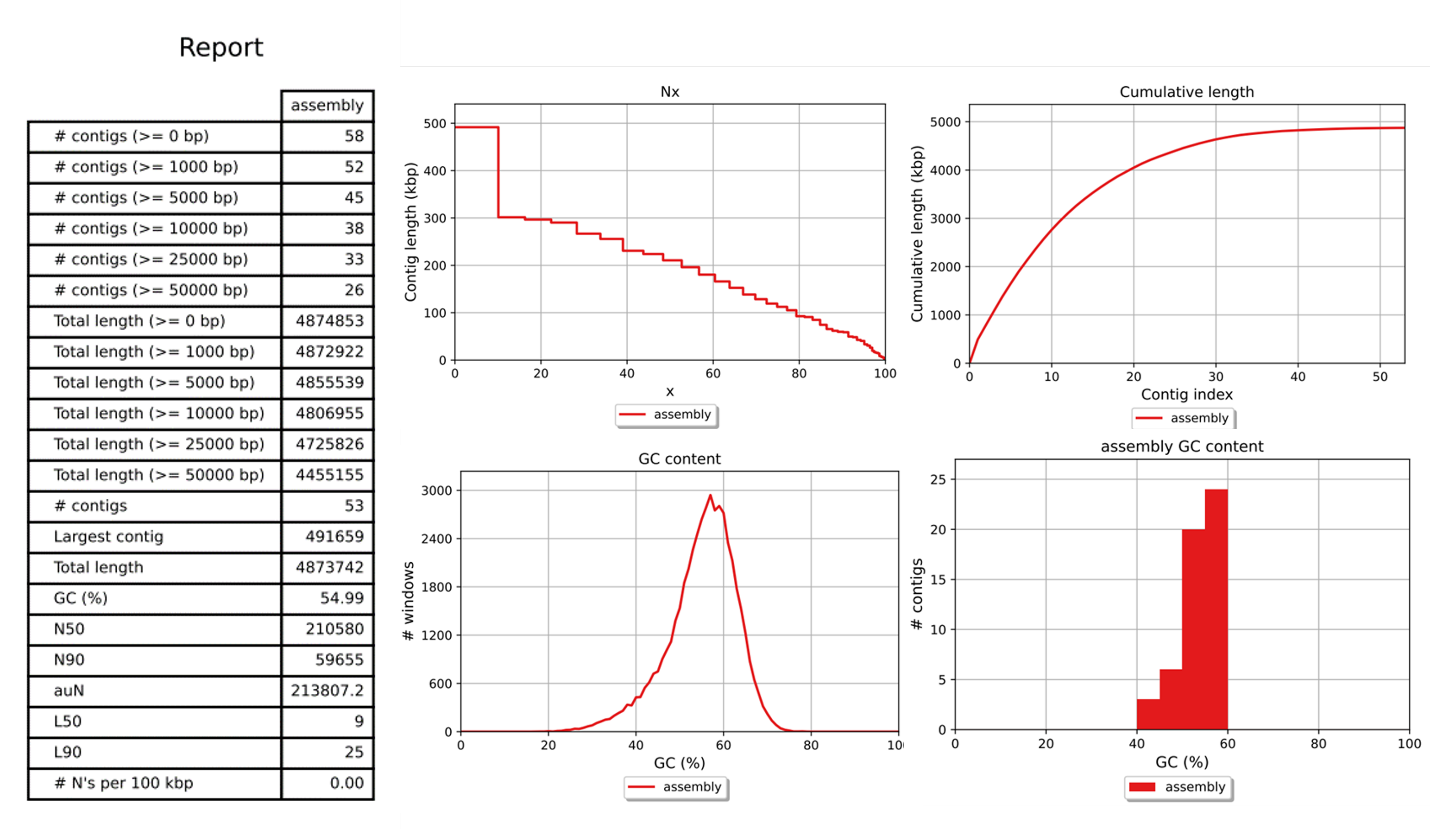

**B8: Assembly quality of the *Proteus mirabilis***

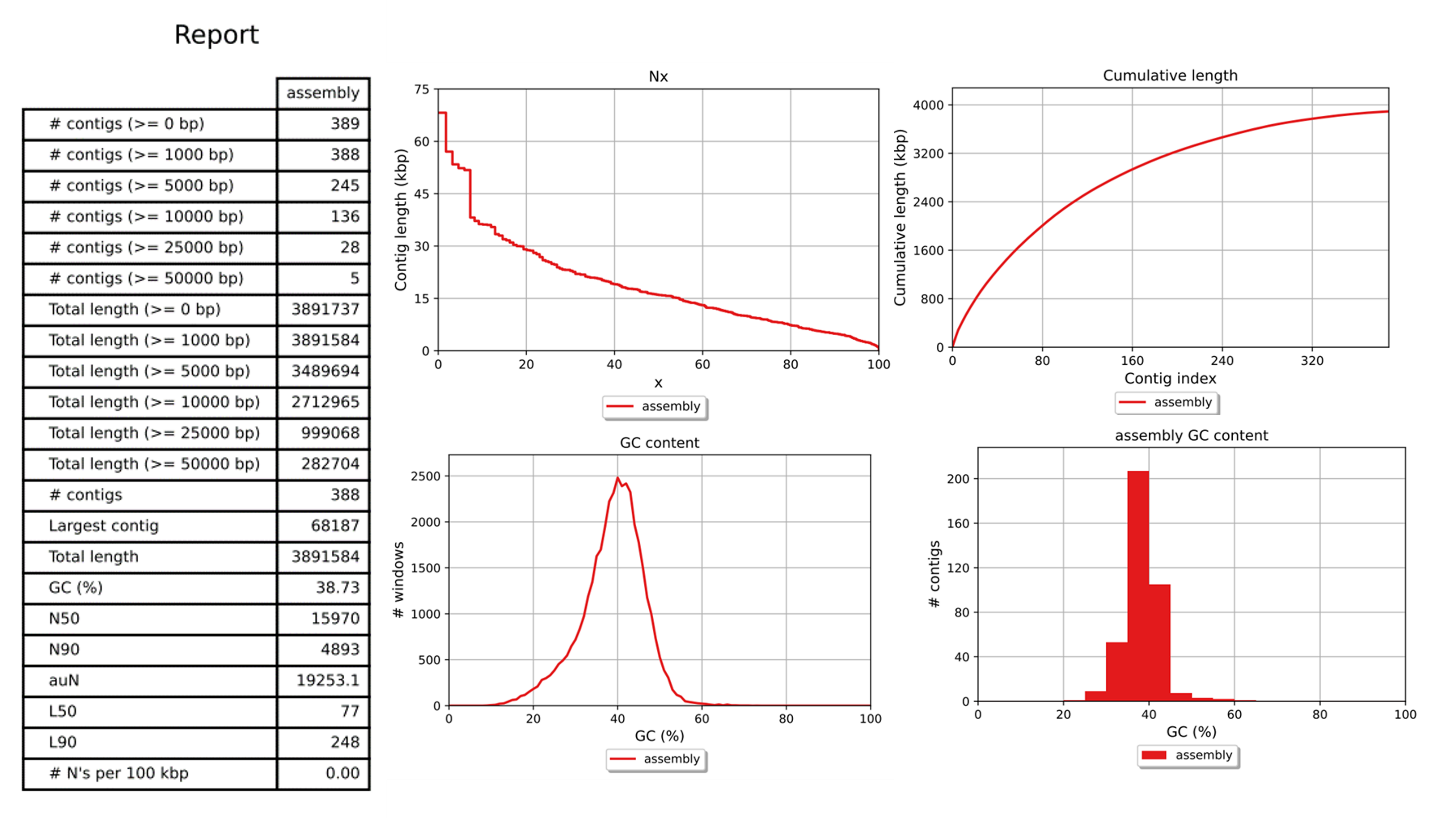

**B9: Assembly quality of the *Citrobacter werkmanii***

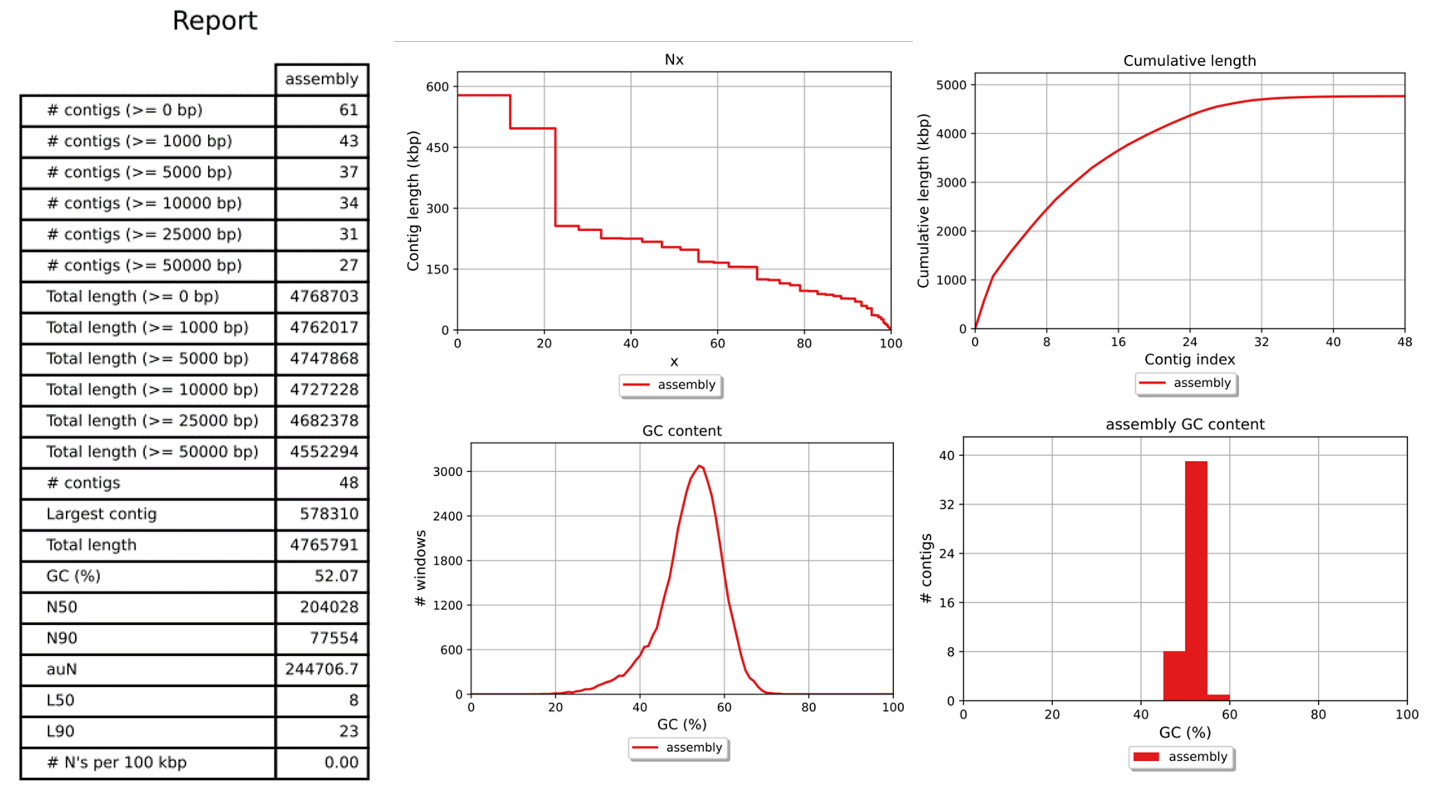

**B10: Assembly quality of the *Providencia stuartii***

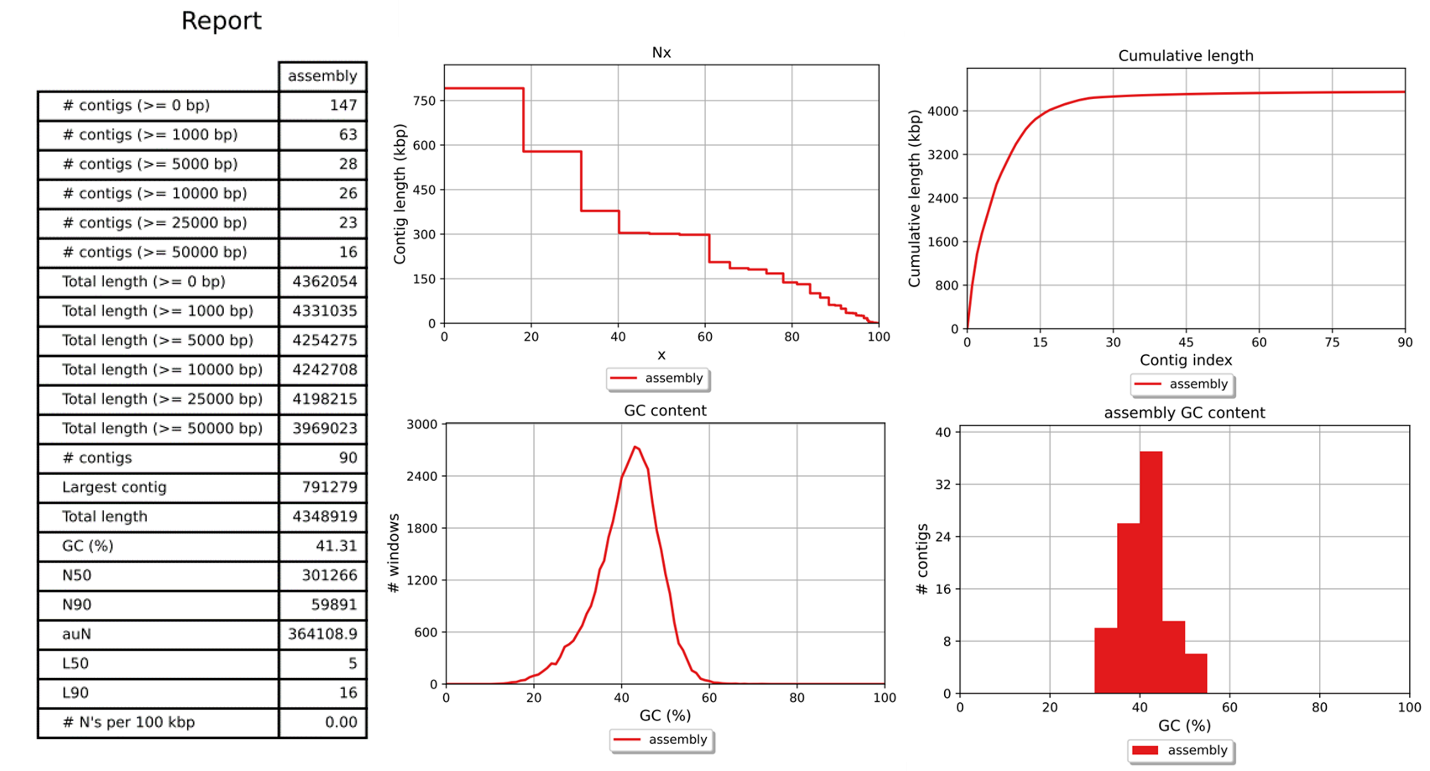
