## Supplementary material for "Linking Genomic Landscape to Disease Mechanism: Core Genetic Factors Underlying Pathogenesis and Antimicrobial Resistance in Diarrheal Pathogens": S5. Virulence Genes

| **Strain Number** | **Bacteria Name** | **Virulence Genes** |
| --- | --- | --- |
| NIB001 | *Plesiomonas shigelloids* | vipB, hcp1 |
| NIB002 | *Providencia stuartii* | **Not found any virulence gene** |
| NIB003 | *Citrobacter werkmanii* | fepC, fepG, entE, entB, entA, fur, gndA, pmrA, ompA, hcp2/tssD2, cgsG, csgE, csgD, csgB, phoP, rcsB, rpoS, allB, acrB |
| NIB006 | *Morganella morganii* | **Not found any virulence gene** |
| NIB007 | *Enterobacter hormaechei* | ompA |
| NIB008 | *Escherichia coli* | ompA, entD, fepA, fes, entF, fepC, fepG, fepD, entS, fepB, entC, entE, entB, entA, espR1, csgB, espX4, espX1, espY1, espY2, espL1, aslA, gspM, gspL, gspK, gspJ, gspI, gspH, gspG, gspF, gspE, gspD, gspC, fimF, fimG, fimH, espX5, espL4, csgD, csgF, csgG, ykgK/ecpR, yagZ/ecpA, yagY/ecpB, yagX/ecpC, yagW/ecpD, yagV/ecpE, fimB, fdeC, fimC |
| NIB009 | *Escherichia coli* | espR4, ybtS, ybtX, ybtQ, ybtP, ybtA, irp2, irp1, ybtU, ybtT, ybtE, fyuA, aslA, espX4, entD, fepA, fes, entF, fepC, fepG, fepD, entS, fepB, entC, entE, entB, entA, espY4, fdeC, gspC, gspD, gspE, gspF, gspG, gspH, gspI, gspJ, gspK, gspL, gspM, kpsM, espL1, espR1, iucA, iucB, iucC, iucD, iutA, espY2, espY1, espX1, ykgK/ecpR, yagZ/ecpA, yagY/ecpB, yagX/ecpC, yagW/ecpD, yagV/ecpE, kpsD, afaE-V, draP, draD, afaC-I, afaB-I, afaA, daaF, ompA, csgG, csgF, csgD, csgB, chuS, shuA, chuT, chuW, shuX, chuY, chuU, chuV, papX |
| NIB010 | *Escherichia coli* | **Not found any virulence gene** |
| NIB011 | *Escherichia coli* | csgB, csgD, csgF, csgG, aslA, chuS, chuA, chuT, chuW, chuX, chuY, chuU, chuV, ybtS, ybtX, ybtQ, ybtP, ybtA, gspL, gspM, kpsM, kpsD, fyuA, ybtE, ybtT, ybtU, irp1, irp2, iroE, iroD, sat, iutA, iucD, iucC, iucB, iucA, fimH, fimG, fimF, fimD, fimC, fimI, fimA, fimE, fimB, afaA, afaB-I, afaC-I, afaD, draP, entD, fepA, fes, entF, fepC, fepG, fepD, entS, fepB, entC, entE, entB, entA, daaF, yagV/ecpE, yagW/ecpD, yagX/ecpC, yagY/ecpB, yagZ/ecpA, ykgK/ecpR, fdeC, papX, ompA |
| NIB012 | *Klebsiella pneumoniae* | entA, entB, fepC, ykgK/ecpR, yagZ/ecpA, yagY/ecpB, yagX/ecpC, yagW/ecpD, yagV/ecpE, ompA |
| NIB013 | *Enterobacter hormaechei* | rcsB, gndA, galF, rpoS, entA, entB, entE, entS, fepD, fepG, fepC, fepA, acrB, phoP, cgsD, ompA, fur, vipA/tssB, vipB/tssC, hcp/tssD, tssF |
| NIB014 | *Klebsiella pneumoniae* | iroE, vipA/tssB, icmF/tssM, tssF, tssG, sciN/tssJ, fur, clpV/tssH, hcp/tssD, ompA, dotU/tssL, vasE/tssK, vipB/tssC, vipA/tssB, entS, fepD, fepG, fepC, entF, fes, fepA, hcp/tssD, fimK, fimH, fimG, fimF, fimD, fimC, fimI, fimA, fimE, fimB, mrkA, mrkB, mrkC, mrkD, mrkF, mrkJ, mrkI, mrkH, rpoS, fepB, entC, entE, entB, entA, ompA, iutA, ykgK/ecpR, yagZ/ecpA, yagY/ecpB,  yagX/ecpC, yagW/ecpD, yagV/ecpE, acrB, acrA, rcsA, rcsB, galF, KP1_RS17355, KP1_RS17345, gndA, ugd, rfbA, rfbB, KP1_RS17240, rfbD, KP1_RS17230, KP1_RS17225, KP1_RS17220 |
| NIB015 | *Escherichia coli* | gspM, gspL, gspK, gspJ, gspI, gspH, gspG, gspF, gspE, gspD, gspC, espL1, entD, fepA, fes, entF, fepC, fepG, fepD, entS, fepB, entC, entE, entB, entA, aslA, yagV/ecpE, yagW/ecpD, yagX/ecpC, yagY/ecpB, yagZ/ecpA, ykgK/ecpR, fdeC, ompA, csgB, csgD, csgF, csgG, espR1, espX5, espX4, espL4, iucA, iucB, iucC, iucD, iutA, espR4, ybtS, ybtX, ybtQ, ybtP, ybtA, irp2, irp1, ybtU, ybtT, ybtE, fyuA, espX1, espY1 |
| NIB018 | *Escherichia fergusonii* | fur, entA, entB, entE, entC, fepB, entS, fepD, fepG, fepC, entF, fes, fepA, entD, ibeB, acrB, gspC, gspD, gspE, gspF, gspG, gspH, gspI, gspJ, gspK, gspL, gspM, gndA, aslA, pmrA, ompA, rcsB, ibeC, allB, csgA, csgB, csgD, cgsE, cgsF, cgsG, tssA, tssM, tssM, tssA, clpV/tssH, tssL, tssJ, hcp2/tssD2, rpoS |
| NIB020 | *Escherichia coli* | rcsB, espR1, csgB, csgA, csgC, phoP, ibeC, rpoS, fur, entA, entB, entE, entC, fepB, entS, fepD, fepG, fepC, fepE, entF, fes, fepA, entD, espX1, galF, KP1_RS17355, espX5, acrB, ompA, pmrA, gspM, gspL, gspK, gspJ, gspI, gspH, gspG, gspF, gspE, gspD, gspC, cfaA, cfaB, cfaC, cfaD/cfaE, allB, KP1_RS17280, rfbK1, hcp1/tssD1, espL1, fimF, fimG, fimH, KP1_RS17345, gndA, yagV/ecpE, yagW/ecpD, yagX/ecpC, yagY/ecpB, yagZ/ecpA, ykgK/ecpR, fdeC, ibeB, cgsE, cgsF, cgsG |
| NIB021 | *Escherichia coli* | espX2, ompA, csgG, csgF, csgD, csgB, fimB, fimE, fimA, fimI, fimC, fimD, fimF, fimG, fimH, espX5, espX4, entA, entB, entE, entC, fepB, entS, fepD, fepG, fepC, entF, fes, fepA, entD, espY4, aslA, yagV/ecpE, yagW/ecpD, yagX/ecpC, yagY/ecpB, yagZ/ecpA, ykgK/ecpR, fdeC, shuS, shuA, shuT, chuW, shuX, chuY, chuU, chuV, espR1, papB, papI, gspM, gspL, gspK, gspJ, gspI, gspH, gspG, gspF, gspE, gspD, gspC, espL1, astA, espY2, espY1, espX1, espY3 |
| NIB022 | *Escherichia coli* | csgB, csgD, csgF, csgG, aslA, fyuA, ybtE, ybtT, ybtU, irp1, irp2, ybtA, ybtP, ybtQ, ybtX, ybtS, ompA, gspL, gspM, kpsM, kpsD, iroD, iroE, fimH, fimG, fimF, fimD, fimC, fimI, fimA, fimE, fimB, iucA, iucB, iucC, iucD, iutA, sat, draP, afaD, afaC-I, afaB-I, afaA, chuV, chuU, chuY, chuX, chuW, chuT, chuA, chuS, daaF, entD, fepA, fes, entF, fepC, fepG, fepD, entS, fepB, entC, entE, entB, entA, yagV/ecpE, yagW/ecpD, yagX/ecpC, yagY/ecpB, yagZ/ecpA, ykgK/ecpR, fdeC, papX |
| NIB023 | *Escherichia coli* | kpsD, kpsT, kpsM, gspM, gspL, gspK, gspJ, gspI, gspH, gspG, gspF, gspE, gspD, gspC, shuS, shuA, shuT, chuW, shuX, chuY, chuU, chuV, espY4, espX5, espX4, espR1, espX1, espY1, espY2, espR4, aslA, espL1, yagV/ecpE, yagW/ecpD, yagX/ecpC, yagY/ecpB, yagZ/ecpA, ykgK/ecpR, fdeC, fyuA, ybtE, ybtT, ybtU, irp1, irp2, ybtA, ybtP, ybtQ, ybtX, ybtS, astA, agg3A, agg3B, agg3C, agg3D, espL4, fimH, fimG, fimF, fimD, fimC, fimI, fimA, fimE, fimB, aap/aspU, entA, entB, entE, entC, fepB, entS, fepD, fepG, fepC, entF, fes, fepA, entD, ompA, csgG, csgF, csgD, csgB, espY3 |
| NIB024 | *Escherichia coli* | espR3, espR4, iucA, iucB, iucC, iucD, iutA, espY4, csgG, espL1, fimG, fimF, aslA, entD, fepA, fes, entF, fepC, fepG, fepD, entS, fepB, entC, entE, entB, entA, espX1, espY1, ompA, chuV, chuU, chuY, shuX, chuW, shuT, shuA, shuS, ybtS, ybtX, ybtQ, ybtP, ybtA, irp2, irp1, ybtU, ybtT, ybtE, fyuA, espL4, espY3, espY2, csgB, fimB, fimE, fimA, fimI, fimC |
| NIB025 | *Proteus mirabilis* | **Not found any virulence gene** |
| NIB026 | *Escherichia coli* | yagV/ecpE, yagW/ecpD, yagX/ecpC, yagY/ecpB, yagZ/ecpA, ykgK/ecpR, fdeC, espR4, espX5, espX4, entD, fepA, fes, entF, fepC, fepG, fepD, entS, fepB, entC, entE, entB, entA, ompA, espL1, espR1, gspC, gspD, gspE, gspF, gspG, gspH, gspI, gspJ, gspK, gspL, gspM, espX1, fimH, fimG, fimF, fimD, fimC, fimI, fimA, fimE, fimB, csgB, csgD, csgF, csgG |
| NIB027 | *Escherichia coli* | entA, entB, entE, entC, fepB, entS, fepD, fepG, fepC, entF, fes, fepA, entD, fdeC, ykgK/ecpR, yagZ/ecpA, yagY/ecpB, yagX/ecpC, yagW/ecpD, yagV/ecpE, ybtA, ybtP, ybtQ, ybtX, ybtS, aslA, chuS, chuA, chuT, chuW, chuX, chuY, chuU, chuV, fyuA, ybtE, ybtT, ybtU, irp1, irp2, sat, iutA, iucD, iucC, iucB, iucA, kpsD, kpsM, gspM, gspL, fimB, fimE, fimA, fimI, fimC, fimD, fimF, fimG, fimH, draP, afaD, afaC-I, afaB-I, afaA, daaF, csgB, csgD, csgF, csgG, ompA, papB, papI, papX |
| NIB029 | *Enterobacter chuandaensis* | acrB, acrA, fepA, fepC, fepD, entS, entE, entB, entA, fur, ompA, csgG, cgsE, phoP, rpoS, tssF, rcsB, galF, gndA, tssF, hcp/tssD, dotU/tssL, hcp/tssD |
| NIB031 | *Escherichia coli* | espR1, espL1, espR3, csgB, csgD, csgF, csgG, ompA, espY4, aslA, entA, entB, entE, entC, fepB, entS |
| NIB032 | *Klebsiella pneumoniae* | fepC, entB, entA, ompA, yagV/ecpE, yagW/ecpD, yagX/ecpC, yagY/ecpB, yagZ/ecpA, ykgK/ecpR |
| NIB033 | *Escherichia coli* | fyuA, ybtE, ybtT, ybtU, irp1, irp2, ybtA, ybtP, ybtQ, ybtX, ybtS, aslA, gspC, gspD, gspE, gspF |
| NIB034 | *Escherichia coli* | ompA, espX4, espX5, gspM, gspL, gspD, gspC, entD, fepA, fes, entF, fepC, fepG, fepD, entS, fepB, entC, entE, entB, entA, espR1, fimB, fimE, fimA, fimI, fimC, fimD, fimF, fimG, fimH, espL1, aslA, yagV/ecpE, yagW/ecpD, yagX/ecpC, yagY/ecpB, yagZ/ecpA, ykgK/ecpR, espX1, csgG, csgF, csgD, csgB |
| NIB035 | *Escherichia coli* | espX1, espY1, espY2, aslA, fyuA, ybtE, ybtT, ybtU, irp1, irp2, ybtA, ybtP, ybtQ, ybtX, ybtS, espR4, entD, fepA, fes, entF, fepC, fepG, fepD, entS, fepB, entC, entE, entB, entA, espX4, chuV, chuU, chuY, shuX, chuW, chuT, shuA, chuS, ompA, fdeC, ykgK/ecpR, yagZ/ecpA, yagY/ecpB, yagX/ecpC, yagW/ecpD, yagV/ecpE, espL1, espR1, afaF-III, draA, draB, afaC-I, afaD, draP, iroD, iroE, iucA, iucB, iucC, iucD, iutA, sat, csgG, csgF, csgD, kpsD, kpsM, gspM, gspL, gspK, gspJ, gspI, gspH, gspG, gspF, gspE, gspD, gspC, espY4, papI, papB, csgB, papX |
| NIB036 | *Escherichia coli* | ybtS, ybtX, ybtQ, ybtP, ybtA, aslA, entA, entB, entE, entC, fepB, entS, fepD, fepG, fepC, entF, fes, fepA, entD, gspL, gspM, kpsM, kpsD, irp2, irp1, ybtU, ybtT, ybtE, fyuA, fimH, fimG, fimF, fimD, fimC, fimI, fimA, fimE, fimB, csgB, csgD, csgF, csgG, ompA, iucA, iucB, iucC, iucD, iutA, sat, draP, afaD, afaC-I, afaB-I, daaF, yagV/ecpE, yagW/ecpD, yagX/ecpC, yagY/ecpB, yagZ/ecpA, ykgK/ecpR, fdeC, papI, papB, papX, afaA, chuV, chuU, chuY, chuX, chuW, chuT, chuA, chuS |
| NIB037 | *Klebsiella pneumoniae* | ykgK/ecpR, yagZ/ecpA, yagY/ecpB, yagX/ecpC, yagW/ecpD, yagV/ecpE, ompA, entA, entB, fepC, fyuA, ybtE, ybtT, ybtU, irp1, irp2, ybtA, ybtP, ybtQ, ybtX, ybtS |
| NIB044 | *Escherichia coli* | ompA, csgB, csgD, csgF, csgG, espL1, espX5, espX4, espR1, entA, entB, entE, entC, fepB, entS, fepD, fepG, fepC, entF, fes, fepA, entD, ybtS, ybtX, ybtQ, ybtP, ybtA, irp2, irp1, ybtU, ybtT, ybtE, fyuA, iutA, espX1, iucA, iucB, iucC, iucD, iutA, yagV/ecpE, yagW/ecpD, yagX/ecpC, yagY/ecpB, yagZ/ecpA, ykgK/ecpR, fdeC, iucB, iucA, afaF-III, draA, draB, draC, draD, draP, draE |
