## Supplementary material for "Linking Genomic Landscape to Disease Mechanism: Core Genetic Factors Underlying Pathogenesis and Antimicrobial Resistance in Diarrheal Pathogens": S9. Genomic Assembly and Annotation Metrics

**Genomic Assembly and Annotation Metrics of the Ten Bacterial Isolates**

| **Bacteria name** | **Strain** | **Host** | **Isolation source** | **Genome size** | **Number of contig** | **Contig N50** | **GC%** | **CDS** | **rRNA** | **tRNA** | **tmRNA** |
| --- | --- | --- | --- | --- | --- | --- | --- | --- | --- | --- | --- |
| ***E. coli*** | NIB044 | Homo sapiens | stool | 4.6 Mb | 166 | 81.1 Kb | 50.75 | 4400 | 4 | 74 | 1 |
|  | NIB015 | Homo sapiens | stool | 4.9 Mb | 119 | 142.8 Kb | 50.71 | 4564 | 4 | 81 | 1 |
|  | NIB008 | Homo sapiens | stool | 4.5 Mb | 109 | 103 Kb | 50.84 | 4216 | 5 | 78 | 1 |
|  | NIB009 | Homo sapiens | stool | 5.4 Mb | 109 | 196.6 Kb | 50.35 | 5098 | 3 | 86 | 1 |
|  | NIB010 | Homo sapiens | stool | 4.8 Mb | 143 | 62.6 Kb | 50.9 | 4490 | 5 | 83 | 1 |
|  | NIB011 | Homo sapiens | stool | 5 Mb | 83 | 169.6 Kb | 50.65 | 4733 | 3 | 78 | 1 |
|  | NIB020 | Homo sapiens | stool | 4.5 Mb | 167 | 69.2 Kb | 50.75 | 4216 | 3 | 45 | 1 |
|  | NIB021 | Homo sapiens | stool | 5.2 Mb | 170 | 140.5 Kb | 50.48 | 4742 | 2 | 74 | 1 |
|  | NIB022 | Homo sapiens | stool | 5.1 Mb | 90 | 169.6 Kb | 50.68 | 4779 | 3 | 77 | 1 |
|  | NIB023 | Homo sapiens | stool | 5.1 Mb | 95 | 127 Kb | 50.5 | 4688 | 5 | 86 | 1 |
|  | NIB024 | Homo sapiens | stool | 4.8 Mb | 225 | 626.1 Kb | 50.83 | 4486 | 3 | 60 | 1 |
|  | NIB026 | Homo sapiens | stool | 4.9 Mb | 145 | 165.3 Kb | 50.74 | 4587 | 3 | 78 | 1 |
|  | NIB027 | Homo sapiens | stool | 5 Mb | 94 | 216.7 Kb | 50.62 | 4756 | 4 | 80 | 1 |
|  | NIB031 | Homo sapiens | stool | 5.2 Mb | 131 | 159.9 Kb | 50.51 | 4869 | 4 | 80 | 1 |
|  | NIB033 | Homo sapiens | stool | 5 Mb | 174 | 163.2 Kb | 50.7 | 4679 | 3 | 79 | 1 |
|  | NIB034 | Homo sapiens | stool | 4.7 Mb | 116 | 148.1 Kb | 50.68 | 4397 | 4 | 71 | 1 |
|  | NIB035 | Homo sapiens | stool | 5.3 Mb | 211 | 196.9 Kb | 50.48 | 4902 | 4 | 83 | 1 |
|  | NIB036 | Homo sapiens | stool | 5.1 Mb | 167 | 172.6 Kb | 50.62 | 4812 | 3 | 76 | 1 |
| ***E. hormaechei*** | NIB013 | Homo sapiens | stool | 4.9 Mb | 56 | 210.6 Kb | 54.99 | 4598 | 4 | 75 | 1 |
|  | NIB007 | Homo sapiens | stool | 4.6 Mb | 73 | 114 Kb | 55.24 | 4390 | 5 | 76 | 1 |
| ***P. mirabilis*** | NIB025 | Homo sapiens | stool | 3.9 Mb | 388 | 15.9 Kb | 38.73 | 3394 | 4 | 54 | 1 |
| ***K. pneumoniae*** | NIB012 | Homo sapiens | Stool | 5.3 Mb | 92 | 242.1 Kb | 57.36 | 4945 | 6 | 64 | 1 |
|  | NIB014 | Homo sapiens | Stool | 5.4 Mb | 63 | 212.7 Kb | 57.39 | 5057 | 4 | 78 | 1 |
|  | NIB032 | Homo sapiens | Stool | 5.3 Mb | 92 | 142.8 Kb | 57.42 | 4973 | 3 | 68 | 1 |
|  | NIB037 | Homo sapiens | Stool | 5.7 Mb | 141 | 183.4 Kb | 57.06 | 5303 | 6 | 65 | 1 |
| ***E. chuandaensis*** | NIB029 | Homo sapiens | Stool | 4.7 Mb | 17 | 672.6 Kb | 55.62 | 4376 | 3 | 74 | 1 |
| ***M. morganii*** | NIB006 | Homo sapiens | Stool | 3.7 Mb | 8 | 2.6 Mb | 51.16 | 3446 | 3 | 67 | 1 |
| ***E. fergusonii*** | NIB018 | Homo sapiens | Stool | 4.5 Mb | 37 | 516.5 Kb | 49.93 | 4220 | 4 | 80 | 1 |
| ***P. stuartii*** | NIB002 | Homo sapiens | stool | 4.3 Mb | 51 | 301.2 Kb | 41.31 | 4006 | 3 | 69 | 1 |
| ***C. werkmanii*** | NIB003 | Homo sapiens | stool | 4.8 Mb | 40 | 204 Kb | 52.07 | 4465 | 4 | 71 | 1 |
| ***P. shigelloides*** | NIB001 | Homo sapiens | stool | 3.6 Mb | 35 | 291.1 Kb | 51.92 | 3096 | 3 | 94 | 1 |
