## Supplementary material for "Linking Genomic Landscape to Disease Mechanism: Core Genetic Factors Underlying Pathogenesis and Antimicrobial Resistance in Diarrheal Pathogens": S10. List of Genomes for Pan-genome Analysis

**List of genomes used for pan-genomic analysis**

| **Bacteria name** | **Strain** | **Host** | **Isolation source** | **Genome size** | **Number of contig** | **Contig N50** | **GC%** | **CDS** | **rRNA** | **tRNA** | **tmRNA** |
| --- | --- | --- | --- | --- | --- | --- | --- | --- | --- | --- | --- |
| ***E. coli*** | PV0838 | *Homo sapiens* | stool | 5.8 Mb | 3 | 5.8 Mb | 50.5 | 5789 | 22 | 108 | 1 |
|  | 2013C-4225 | *Homo sapiens* | stool | 5.7 Mb | 2 | 5.6 Mb | 50.5 | 5595 | 22 | 98 | 1 |
|  | 2010C-3142 | *Homo sapiens* | stool | 5.6 Mb | 2 | 5.5 Mb | 50.5 | 5367 | 22 | 108 | 1 |
|  | NIB044 | *Homo sapiens* | stool | 4.6 Mb | 166 | 81.1 Kb | 50.75 | 4400 | 4 | 74 | 1 |
|  | NIB015 | *Homo sapiens* | stool | 4.9 Mb | 119 | 142.9 Kb | 50.71 | 4564 | 4 | 81 | 1 |
|  | NIB008 | *Homo sapiens* | stool | 4.5 Mb | 109 | 103 Kb | 50.84 | 4216 | 5 | 78 | 1 |
|  | NIB009 | *Homo sapiens* | stool | 5.4 Mb | 109 | 196.6 Kb | 50.35 | 5098 | 3 | 86 | 1 |
|  | NIB010 | *Homo sapiens* | stool | 4.8 Mb | 143 | 62.7 Kb | 50.9 | 4490 | 5 | 83 | 1 |
|  | NIB011 | *Homo sapiens* | stool | 5 Mb | 83 | 169.6 Kb | 50.65 | 4733 | 3 | 78 | 1 |
|  | NIB020 | *Homo sapiens* | stool | 4.5 Mb | 167 | 69.3 Kb | 50.75 | 4216 | 3 | 45 | 1 |
|  | NIB021 | *Homo sapiens* | stool | 5.2 Mb | 170 | 140.6 Kb | 50.48 | 4742 | 2 | 74 | 1 |
|  | NIB022 | *Homo sapiens* | stool | 5.1 Mb | 90 | 169.6 Kb | 50.68 | 4779 | 3 | 77 | 1 |
|  | NIB023 | *Homo sapiens* | stool | 5 Mb | 95 | 126.9 Kb | 50.5 | 4688 | 5 | 86 | 1 |
|  | NIB024 | *Homo sapiens* | stool | 4.8 Mb | 225 | 62.6 Kb | 50.83 | 4486 | 3 | 60 | 1 |
|  | NIB026 | *Homo sapiens* | stool | 4.9 Mb | 145 | 165.3 Kb | 50.74 | 4587 | 3 | 78 | 1 |
|  | NIB027 | *Homo sapiens* | stool | 5 Mb | 94 | 216.7 Kb | 50.62 | 4756 | 4 | 80 | 1 |
|  | NIB031 | *Homo sapiens* | stool | 5. Mb | 131 | 159.9 Kb | 50.51 | 4869 | 4 | 80 | 1 |
|  | NIB033 | *Homo sapiens* | stool | 5 Mb | 174 | 163.2 Kb | 50.7 | 4679 | 3 | 79 | 1 |
|  | NIB034 | *Homo sapiens* | stool | 4.7 Mb | 116 | 148.1 Kb | 50.68 | 4397 | 4 | 71 | 1 |
|  | NIB035 | *Homo sapiens* | stool | 5.3 Mb | 211 | 196.9 Kb | 50.48 | 4902 | 4 | 83 | 1 |
|  | NIB036 | *Homo sapiens* | stool | 5. 2Mb | 167 | 172.5 Kb | 50.62 | 4812 | 3 | 76 | 1 |
| ***E. hormaechei*** | L51 | *Homo sapiens* | feces | 5.4 Mb | 2 | 5 Mb | 54.5 | 5121 | 25 | 86 | 1 |
|  | FY-1 | *Homo sapiens* | blood | 5.2 Mb | 4 | 5 Mb | 55 | 4936 | 25 | 87 | 1 |
|  | A1 | *Homo sapiens* | Rectal swab | 5.4 Mb | 3 | 4.9 Mb | 55 | 5079 | 25 | 86 | 1 |
|  | EB6 | *Homo sapiens* | urine | 5.3 Mb | 3 | 4.9 Mb | 54.5 | 5052 | 25 | 86 | 1 |
|  | C45 | *Homo sapiens* | clinical sample | 5.5 Mb | 5 | 5.1 Mb | 55 | 5259 | 25 | 84 | 1 |
|  | C44 | *Homo sapiens* | clinical sample | 5 Mb | 5 | 4.7 Mb | 55.5 | 4623 | 25 | 86 | 1 |
|  | YY1 | *Homo sapiens* | secretion | 5 Mb | 2 | 4.7 Mb | 55 | 4723 | 25 | 86 | 1 |
|  | C4 | *Homo sapiens* | clinical sample | 5 Mb | 5 | 4.7 Mb | 55.5 | 4635 | 25 | 86 | 1 |
|  | Y-2 | *Homo sapiens* | urine | 4.7 Mb | 4 | 4.7 Mb | 55.5 | 4331 | 25 | 85 | 1 |
|  | Ec61 | *Homo sapiens* | Transtracheal aspirate | 5.1 Mb | 4 | 4.9 Mb | 55 | 4850 | 25 | 86 | 1 |
|  | va18651 | *Homo sapiens* | cervical swab | 5.4 Mb | 5 | 4.8 Mb | 55 | 5118 | 25 | 87 | 1 |
|  | NIB013 | *Homo sapiens* | stool | 4.9 Mb | 56 | 210.6 Kb | 54.99 | 4598 | 4 | 75 | 1 |
|  | NIB007 | *Homo sapiens* | stool | 4.6 Mb | 73 | 114 Kb | 55.24 | 4390 | 5 | 76 | 1 |
| ***P. mirabilis*** | PM52260 | *Homo sapiens* | sputum | 4.3 Mb | 3 | 4.2 Mb | 39.5 | 3934 | 22 | 84 | 1 |
|  | PM52808 | *Homo sapiens* | sputum | 4.3 Mb | 4 | 4.2 Mb | 39.5 | 3935 | 24 | 85 | 1 |
|  | HURS-181823 | *Homo sapiens* | urine | 4.3 Mb | 1 | 4.3 Mb | 39 | 3937 | 22 | 86 | 1 |
|  | HURS-186083 | *Homo sapiens* | Rectal swab | 4.3 Mb | 1 | 4.3 Mb | 39 | 3936 | 22 | 86 | 1 |
|  | P13 | *Homo sapiens* | not applicable | 4.3 Mb | 2 | 4.3 Mb | 39 | 3942 | 22 | 84 | 1 |
|  | P3 | *Homo sapiens* | not applicable | 4.4 Mb | 3 | 4.3 Mb | 39.5 | 4089 | 22 | 84 | 1 |
|  | PM74 | *Homo sapiens* | urine | 4.3 Mb | 2 | 4.2 Mb | 39 | 3897 | 22 | 84 | 1 |
|  | PM33 | *Homo sapiens* | sputum | 4.6 Mb | 3 | 4.3 Mb | 39.5 | 4292 | 22 | 84 | 1 |
|  | PM8762 | *Homo sapiens* | urine | 4.3 Mb | 1 | 4.3 Mb | 39.5 | 3876 | 22 | 84 | 1 |
|  | AOUC-001 | *Homo sapiens* | blood | 4.3 Mb | 1 | 4.3 Mb | 39.5 | 3952 | 22 | 84 | 1 |
|  | NIB025 | *Homo sapiens* | stool | 3.9 Mb | 388 | 15970 | 38.73 | 3394 | 4 | 54 | 1 |
| ***K. pneumoniae*** | KP64 | *Homo sapiens* | human urine | 5.7 Mb | 3 | 5.6 Mb | 57 | 5275 | 25 | 85 | 1 |
|  | KP29105 | *Homo sapiens* | sputum | 5.8 Mb | 3 | 5.5 Mb | 57 | 5357 | 25 | 87 | 1 |
|  | QD23 | *Homo sapiens* | urine | 5.8 Mb | 1 | 5.8 Mb | 57 | 5510 | 25 | 90 | 1 |
|  | Bio45 | *Homo sapiens* | skin swab | 6.1 Mb | 2 | 5.7 Mb | 56 | 5706 | 3 | 73 | 1 |
|  | Bio73 | *Homo sapiens* | sputum of human | 6.1 Mb | 2 | 5.7 Mb | 56.5 | 5701 | 3 | 71 | 1 |
|  | Bio3 | *Homo sapiens* | sputum | 6.1 Mb | 2 | 5.7 Mb | 56.5 | 5688 | 3 | 73 | 1 |
|  | TH12908 | *Homo sapiens* | blood | 5.7 Mb | 1 | 5.7 Mb | 57 | 5239 | 25 | 25 | 1 |
|  | CRE146 | *Homo sapiens* | patient's alveolar lavage fluid | 6.3 Mb | 7 | 5.6 Mb | 56.5 | 6077 | 25 | 88 | 1 |
|  | hvKP340 | *Homo sapiens* | sputum | 6 Mb | 8 | 5.7 Mb | 57 | 5762 | 25 | 85 | 1 |
|  | KP2722 | *Homo sapiens* | sputum | 6.1 Mb | 6 | 5.6 Mb | 57 | 5804 | 25 | 85 | 1 |
|  | NIB012 | *Homo sapiens* | Stool | 5.3 Mb | 92 | 242.1 Kb | 57.36 | 4945 | 6 | 64 | 1 |
|  | NIB014 | *Homo sapiens* | Stool | 5.4 Mb | 63 | 212.6 Kb | 57.39 | 5057 | 4 | 78 | 1 |
|  | NIB032 | *Homo sapiens* | Stool | 5.4 Mb | 92 | 142.7 Kb | 57.42 | 4973 | 3 | 68 | 1 |
|  | NIB037 | *Homo sapiens* | Stool | 5.7 Mb | 141 | 183.4 Kb | 57.06 | 5303 | 6 | 65 | 1 |
| ***E. chuandaensis*** | AEC | *Homo sapiens* | breast milk | 4.9 Mb | 10 | 2.9 Mb | 55.5 | 4560 | 25 | 87 | 1 |
|  | HD8830 | *Homo sapiens* | Surgical wound | 4.8 Mb | 25 | 902.9 kb | 55.5 | 4512 | 3 | 69 | 1 |
|  | M23267 | *Homo sapiens* | urine | 5.1 Mb | 62 | 683.5 kb | 55 | 4820 | 4 | 80 | 1 |
|  | EC9472 | *Homo sapiens* | urine | 4.9 Mb | 48 | 417.4 kb | 55.5 | 4567 | 5 | 73 | 1 |
|  | NIB029 | *Homo sapiens* | Stool | 4.7 Mb | 17 | 672.5 Kb | 55.62 | 4376 | 3 | 74 | 1 |
| ***M. morganii*** | IPS040 | *Homo sapiens* | feces | 3.9 Mb | 35 | 682.5 kb | 51 | 3713 | 3 | 70 | 1 |
|  | CTX51T | *Homo sapiens* | adenocarcinoma in cecum | 4.2 Mb | 2 | 4.2 Mb | 50.5 | 3890 | 22 | 82 | 1 |
|  | MM50821 | *Homo sapiens* | sputum-aspirate | 4.3 Mb | 4 | 4.2 Mb | 51 | 4090 | 22 | 82 | 1 |
|  | MM46903 | *Homo sapiens* | ulcer swab | 4.3 Mb | 4 | 4.1 Mb | 51 | 4007 | 22 | 82 | 1 |
|  | MM48659 | *Homo sapiens* | urine | 4.3 Mb | 3 | 4.2 Mb | 51 | 4025 | 22 | 82 | 1 |
|  | SMM01 | *Homo sapiens* | Midstream Urine | 3.9 Mb | 1 | 3.9 Mb | 51 | 3702 | 19 | 77 | 1 |
|  | L241 | *Homo sapiens* | feces | 3.9 Mb | 2 | 3.9 Mb | 51 | 3635 | 22 | 82 | 1 |
|  | K266 | *Homo sapiens* | sputum | 4.1 Mb | 1 | 4.1 Mb | 51 | 3872 | 22 | 81 | 1 |
|  | NIB006 | *Homo sapiens* | Stool | 3.7 Mb | 8 | 256.9 Kb | 51.16 | 3446 | 3 | 67 | 1 |
| **E. fergusonii** | NIB018 | *Homo sapiens* | Stool | 4.5 Mb | 37 | 516.5 Kb | 49.93 | 4220 | 4 | 80 | 1 |
| ***Providencia stuartii*** | 41 | *Homo sapiens* | Rectal Swab | 4.5 Mb | 2 | 4.3 Mb | 41.5 | 4164 | 22 | 80 | 1 |
|  | CAVP490 | *Homo sapiens* | human | 4.6 Mb | 6 | 4.4 Mb | 41.5 | 4166 | 22 | 78 | 1 |
|  | CAVP450 | *Homo sapiens* | human | 4.7 Mb | 7 | 4.4 Mb | 41.5 | 4199 | 22 | 78 | 1 |
|  | BE2467 | *Homo sapiens* | urine | 4.6 Mb | 3 | 4.4 Mb | 42 | 4316 | 22 | 81 | 1 |
|  | PRV00010 | *Homo sapiens* | sputum | 4.2 Mb | 26 | 679.5 kb | 40.5 | 3738 | 4 | 65 | 1 |
|  | PRV00003 | *Homo sapiens* | urine | 4.5 Mb | 36 | 500.5 kb | 41.5 | 4229 | 8 | 74 | 1 |
|  | PRV00007 | *Homo sapiens* | sputum | 4.3 Mb | 37 | 456.7 kb | 41.5 | 3914 | 10 | 76 | 1 |
|  | PS71 | *Homo sapiens* | blood | 4.4 Mb | 79 | 456 kb | 42 | 4025 | 8 | 77 | 1 |
|  | 3347685 | *Homo sapiens* | decubitus ulcer | 4.7 Mb | 2 | 4.5 Mb | 42 | 4326 | 22 | 81 | 1 |
|  | NIB002 | *Homo sapiens* | stool | 4.3 Mb | 51 | 301.2 Kb | 41.31 | 4006 | 3 | 69 | 1 |
| ***Citrobacter werkmanii*** | FDAARGOS_364 | *Homo sapiens* | stool | 4.9 Mb | 1 | 4.9 Mb | 52.5 | 4571 | 25 | 85 | 1 |
|  | NBRC 105721 | *Homo sapiens* | human blood | 4.9 Mb | 30 | 427.6 kb | 52 | 4633 | 4 | 71 | 1 |
|  | 28_P_CW | *Homo sapiens* | rectal | 5.2 Mb | 3 | 5.1 Mb | 52 | 4983 | 25 | 85 | 1 |
|  | 2023EL00951 | *Homo sapiens* | Rectal Swab | 5.1 Mb | 22 | 2.7 Mb | 52 | 4780 | 10 | 78 | 1 |
|  | CRE806 | *Homo sapiens* | slough from diabetec foot ulcer | 5.3 Mb | 121 | 248.8 kb | 52 | 5045 | 10 | 78 | 1 |
|  | CB00078 | *Homo sapiens* | wound | 5.1 Mb | 77 | 242.5 kb | 50.5 | 4690 | 4 | 73 | 1 |
|  | CRE1173 | *Homo sapiens* | pus | 5.4 Mb | 124 | 218 kb | 52 | 5170 | 9 | 76 | 1 |
|  | K48_1 | *Homo sapiens* | feces | 5.1 Mb | 91 | 206 kb | 52 | 4812 | 5 | 64 | 1 |
|  | CRE1470 | *Homo sapiens* | peritoneal fluid | 5.2 Mb | 105 | 199 kb | 52 | 4898 | 7 | 73 | 1 |
|  | NIB003 | *Homo sapiens* | stool | 4.7 Mb | 40 | 204 Kb | 52.07 | 4465 | 4 | 71 | 1 |
| ***P. shigelloides*** | NCTC10363 | *Homo sapiens* | feces | 3.8 Mb | 1 | 3.5 Mb | 51.5 | 3338 | 37 | 119 | 1 |
|  | S2023588 | *Homo sapiens* | patient with gastroenteritis | 3.5 Mb | 30 | 261.3 kb | 52 | 3160 | 3 | 81 | 1 |
|  | S2023591 | *Homo sapiens* | patient with gastroenteritis | 3.7 Mb | 30 | 230.7 kb | 52 | 3158 | 3 | 88 | 1 |
|  | NCTC10364 | *Homo sapiens* | feces | 3.7 Mb | 38 | 200.5 kb | 52 | 3131 | 44 | 115 | 1 |
|  | S2023589 | *Homo sapiens* | patient with gastroenteritis | 3.7 Mb | 33 | 211.7 kb | 52 | 3158 | 3 | 78 | 1 |
|  | NIB001 | *Homo sapiens* | stool | 3.6 Mb | 35 | 291.1 Kb | 51.93 | 3096 | 3 | 94 | 1 |
